# Design of D-amino acids SARS-CoV-2 Main protease inhibitors using the cationic peptide from rattlesnake venom as a scaffold

**DOI:** 10.1101/2021.11.10.468025

**Authors:** Raphael J. Eberle, Ian Gering, Markus Tusche, Philipp N. Ostermann, Lisa Müller, Ortwin Adams, Heiner Schaal, Danilo S. Olivier, Marcos S. Amaral, Raghuvir K. Arni, Dieter Willbold, Mônika A. Coronado

## Abstract

The C30 Endopeptidase (3C-like protease; 3CL^pro^) is essential for the life cycle of SARS-CoV-2 (severe acute respiratory syndrome-coronavirus-2) since it plays a pivotal role in viral replication and transcription and is hence a promising drug target. Molecules isolated from animals, insects, plants or microorganisms can serve as a scaffold for the design of novel biopharmaceutical products. Crotamine, a small cationic peptide from the venom of the rattlesnake *Crotalus durissus terrificus* has been the focus of many studies since it exhibits activities such as analgesic, in vitro antibacterial and hemolytic activities. The crotamine derivative L-peptides (L-CDP) that inhibit the 3CL protease in the low µM range were examined since they are susceptible to proteolytic degradation; we explored the utility of their D-enantiomers form. Comparative uptake inhibition analysis showed D-CDP as a promising prototype for a D-peptide-based drug. We also found that the D-peptides can impair SARS-CoV-2 replication *in vivo*, probably targeting the viral protease 3CL^pro^.

## 1. Introduction

In Wuhan, Hubei Province, China, in December 2019, a rapid increase in the number of pneumonia suspect cases [1] quickly aroused interest, and it sounded the emergency call in the World Health Organization (WHO) in January 2020, as a “public health emergency of international concern”. The resulting disease, Coronavirus Disease-2019 (COVID-19), exploded into a global pandemic within a few months, claiming lives in all continents. SARS-Coronavirus-2 (SARS-CoV-2) has resulted in over 248 million confirmed cases and over 5 million deaths worldwide reported by the WHO on 5th of November 2021 [2]. The worldwide vaccination campaign using clinical safe and efficient vaccines against SARS-CoV-2 (e.g. BioNTech-Pfizer, Moderna, Johnson & Johnson, and AstraZeneca vaccines) [3-6], and so far, more than 7 billion vaccine doses have been administered [2].

Several candidate drugs that may inhibit SARS-CoV-2 infection and replication have been approved for emergency use (e.g., Remdesivir, dexamethasone, Favipiravir, Lopinavir/ritonavir, and Darunavir) [7-11]. Given a considerable limitation of direct-acting antivirals for COVID-19 and an increasing presence of SARS-CoV-2 variants (B.1.617.2, B.1.1.7, B.1.351, A.23.1, B.1.525, B.1.526 and P.1) [12], it remains a strategic priority to develop new candidates with minimal side effects and which are also targeted against new variants. Upon entering and uncoating the viral particles, the positive-stranded RNA genome is rapidly translated into two polyproteins processed (pp1a and pp1ab) by 3CL and papain-like proteases into 16 nonstructural proteins (NSPs) [13,14]. 3CL^pro^ is a cysteine protease organised in three domains (domains I to III) with a chymotrypsin-like fold [15]. Its active form consists of two protomers (homodimer) containing a noncanonical Cys-His dyad located in the cleft between domains I and II [15-17]. The functional importance of 3CL^pro^ in the viral life cycle combined with the absence of closely related homologues in humans indicates that this protease is an attractive target for developing antiviral drugs [18].

In recent years, antimicrobial peptides (AMPs) have been considered to hold the promise as a viable solution to form the basis for the design of novel peptides to combat hazardous microorganism infections. The use of AMPs can be promising as a therapeutic tool to address increasing viral infections, for which no current or authorised medication or treatment is available [19]. AMPs are frequently used to treat viral-related diseases such as Zika (ZIKV), Dengue (DENV) [20], and Influenza A virus infection (IAV) [21].

Crotamine (Cro), a small cationic polypeptide originally encountered in the venom of the South American rattlesnake *Crotalus durissus terrificus* [22, 23], possess cell wall penetrating properties [24,25], and several biological functions of this polypeptide were described, including antimicrobial, antifungal and antitumoral activities [24-27]. These properties were mainly considered to be determined by the overall positive net surface charge distribution of crotamine [24-27]. Characterised as a novel cell-penetrating polypeptide (CPP) nanocarrier, Cro has biotechnological applications due to its peculiar specificity for highly proliferating cells. [24,25,28]. Similar to other CPPs, Cro showed a rapid translocation efficiency (within 5 min) into all cell types investigated to date [29]. A small peptide composed by 42 amino acid residues (YKQCHKKGGHCFPKEKICLPPSSDFGKMDCRWRWKCCKKGSG) [22, 23] containing two putative nuclear localization sequence (NLS) motifs Cro_2-18 (KQCHKKGGHCFPKEKIC) and Cro_27-39 (KMDCRWRWKCCKK) [29]. The sequence Crot_27-39 was selected and named L-CDP1 as the initial sequence for inhibition studies against SARS-CoV-2 3CL^pro^. Based on the first sequence, several Crotamine Derivative Peptides (CDPs) in L-form were specifically modified, and D-forms were designed. They were tested concerning their inhibitory potential against the virus’ main protease.

## 2. Results and Discussion

### 2.1. Preparation of SARS-CoV-2 3CL^pro^

SARS-CoV-2 3CL^pro^_GST fusion protein was expressed in *E. coli* Lemo21 (DE3) cells and purified using a GSH-Sepharose column (Supplementary Fig. S3A). The relevant protein fractions were concentrated and prepared for PreScission protease cleavage to remove the GST-tag. The SDS gel (Supplementary Fig. S1) indicates the cleavage efficiency and the purity of 3CL^pro^.

### 2.2. Primary inhibition assay of Crotamine and L-CDPs against SARS-CoV-2 3CL^pro^

The L-CDP1 (wild type sequence Cro_27-39) includes three cysteine residues. The L-CDP2-9 are modified peptides by substituting the cysteine to serine residues in a different position (Table 1 and Supplementary Fig. S2 and S3) to efficiently achieve an optimised sequence aiming to inhibit the 3CLpro. In principle, the positively charged residues were maintained due to their membrane modify properties.

**Table 1.**
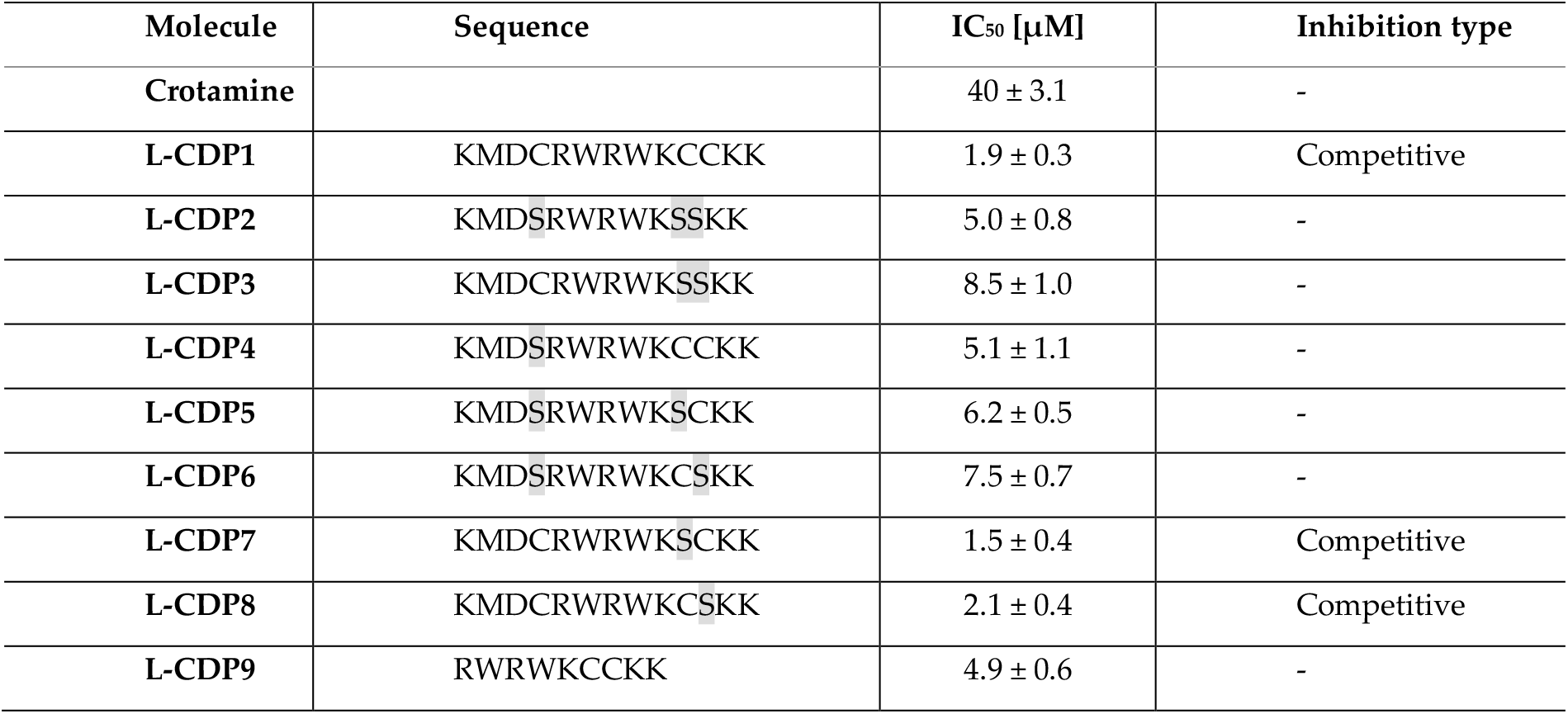
Summary of the SARS-CoV-2 3CLpro inhibition experiments by Crotamine and CDPs.

According to the procedure described earlier, the SARS-CoV-2 3CL^pro^ activity assay was performed using DABCYL-KTSAVLQ/SGFRKME-EDANS (Bachem, Switzerland) as substrate [2-5]. A primary inhibition test with Cro and the L-CDPs (30 µM) was performed to screen the best inhibitor peptide against the virus protease (Fig. 1).

**Figure 1.**
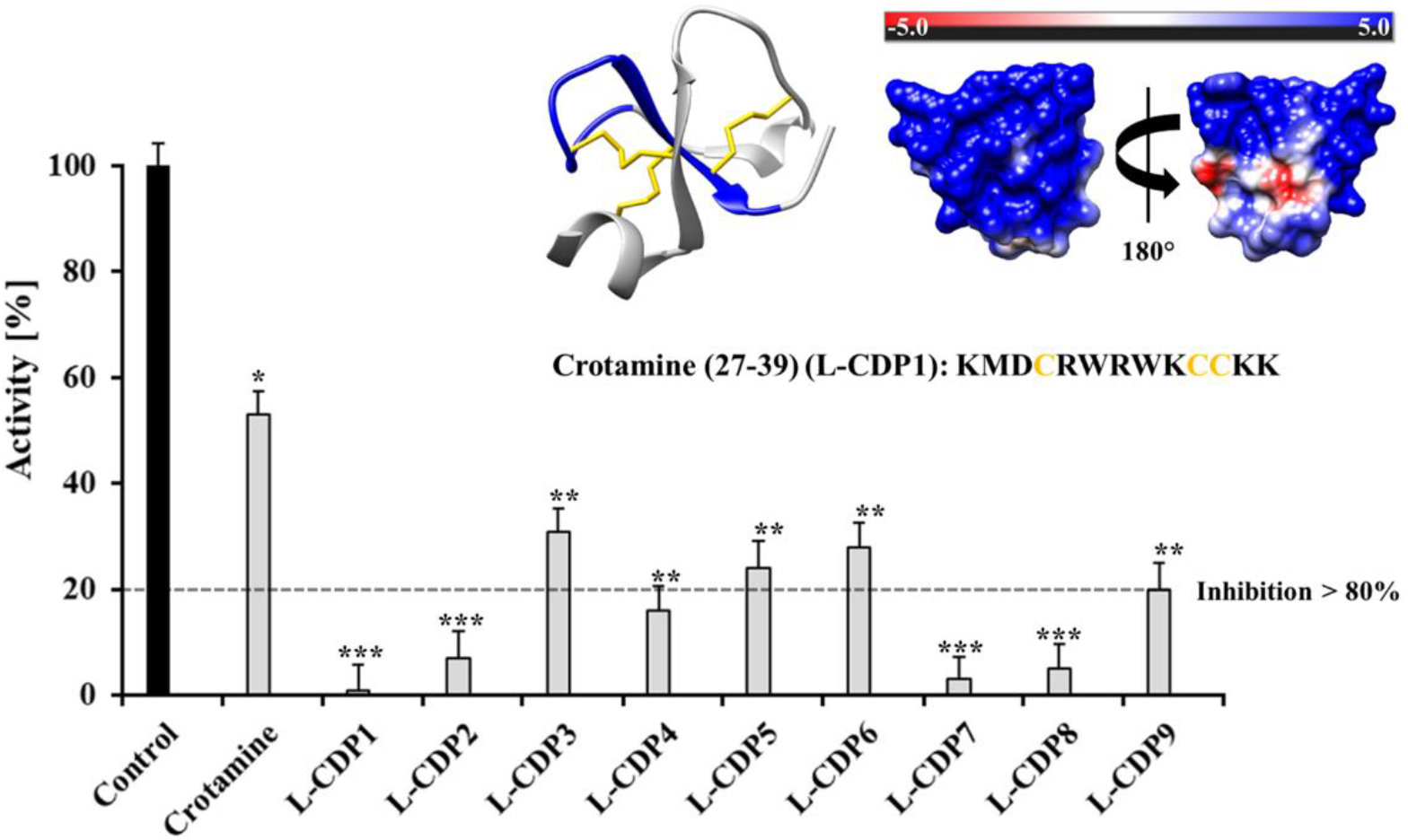
Primary inhibition tests of Crotamine and L-CDPs against SARS-CoV-2 3CLpro. Crotamine inhibits the virus protease activity by around 50%. L-CDP1, L-CDP2, L-CDP4, L-CDP7 and L-CDP8 inhibit the virus protease activity by more than 80%. Data shown are the mean ± SD from 3 independent measurements (n=3). Asterisks mean that the data differs from the control (0 µM inhibitor) significantly at p < 0.05 (*), p < 0.01 (**) and p < 0.001 (***), level according to ANOVA and Tukey’s test. The experimental model of Crotamine is shown in coulombic surfaces and cartoons, with the L-CDP1 sequence highlighted in blue (PDB entry: 4GV5).

The primary inhibition tests revealed a strong effect of L-CDP1, L-CDP2, L-CDP7 and L-CDP8 peptides against SARS-CoV-2 3CL^pro^ activity assay, and those peptides inhibit the virus protease activity by more than 80%. All peptides were subject of the dose-dependent studies (Supplementary Fig. S4); however, the peptides that present more than 80% inhibition have been chosen for further studies.

### 2.3. Characterisation of the 3CLpro inhibition by Crotamine and L-CDPs

The selected peptides were further analysed with respect to their actual potential to inhibit the catalytic activity of the 3CL^pro^ in a biochemical assay. To gain insight into functional implications caused by the selected peptide (L-CDP1), we checked the minimum concentration of Cro required to inhibit 100% of the protease activity. The full-length polypeptide Cro was tested using a concentration range of 0-300 µM, and the polypeptide demonstrates 100% protease inhibition at a concentration of 300 µM (Supplementary Fig. S4A), presenting an IC_50_ value of 40 ± 3.1 µM (Table 1, Fig. 2A). In contrast, the selected wild type sequence Crot_27-39 (L-CDP1) inhibits 100% of the recombinant SARS-CoV-2 protease activity at a concentration of 30 µM (Supplementary Fig. S4B) with IC_50_ value of 1.9 ± 0.3 µM (Table 1, Fig.2B). The replacement of all cysteine residues of L-CDP1 by serine residues, a peptide named L-CDP2 (Table 1), led to an increase in IC_50_ values (5.0 ± 0.8 µM), revealing that the cysteine residue can increase the inhibition rate of the protease (Fig. 2C and Supplementary Fig. S4C). The Crotamine derivative L-peptide-7, a peptide that presents an amino acid substitution, Cys36 (WT numbering) to Ser, and L-CDP8 (Cys37 to Ser) (see Table 1), inhibits the protease by 100% at 60 µM concentration (Supplementary Fig. S4HI) with IC_50_ values of 1.5 ± 0.4 and 2.1 ± 0.4 µM (Table 1, Fig. 2DE), respectively. The Supplementary Fig. S4 shows the inhibition effect for all modified L-CDP against the 3CL protease and the Supplementary Fig. S5 the dose-response curves for IC_50_ determination for the remaining modified peptides (L-CDP3, 4, 5, 6 and 9).

**Figure 2.**
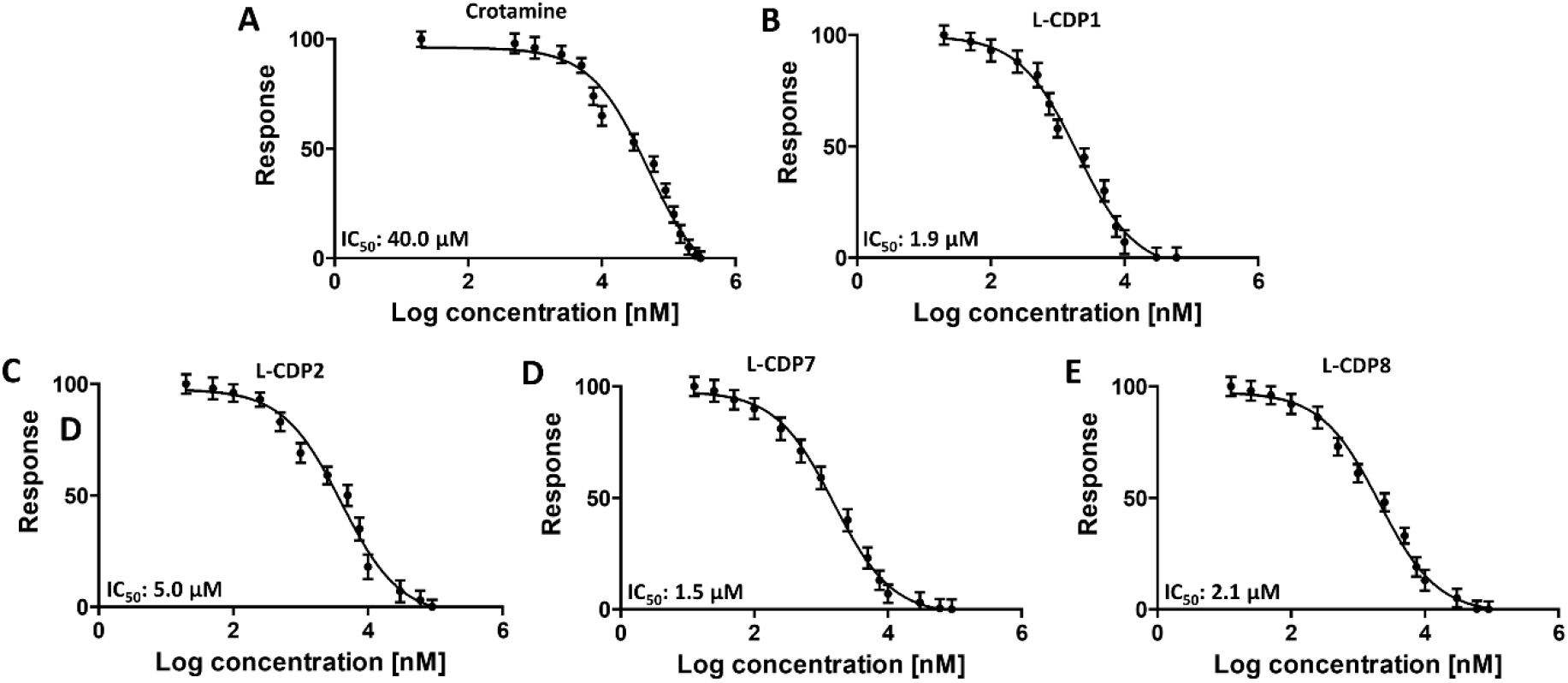
Crotamine, L-CDP1, L-CDP2, L-CDP7 and L-CDP8 inhibitory activity against SARS-CoV-2 3CLpro. Dose-response curves for IC_50_ determination. The normalised response [%] of SARS-CoV-2 3CLpro is plotted against the Log of the inhibitor concentration **A:** Dose-response curve of Crotamine and SARS-CoV-2 3CLpro. **B:** Dose-response curve of L-CDP1 and SARS-CoV-2 3CLpro. **C:** Dose-response curve of L-CDP2 and SARS-CoV-2 3CLpro. **CD** Dose-response curve of L-CDP7 and SARS-CoV-2 3CLpro. **E:** Dose-response curve of L-CDP8 and SARS-CoV-2 3CLpro. Data shown are the mean ± SD from three independent measurements (n=3).

Regarding residues substitution (Table 1), the secondary structure of L-CDP1, L-CDP2, L-CDP7 and L-CDP8 was investigated by Circular dichroism (CD), demonstrating that the substitution of the cysteine residue will results in conformational changes (Supplementary Fig. S6). As expected, difference in uptake behavior was observed with the replacement of only one cysteine residue, as observed by IC50 values (Table 1 and Fig. 2C). However, the substitution at position 36 (WT numbering) increases the inhibitory efficiency of the peptide L-CDP7 (Table 1 and Fig. 2D). Table 1 summarises the experiments performed with the modified L-CDP peptides.

Detailed mechanistic studies using a fluorescence-based protease assay demonstrated that L-CDP1, L-CDP7, and L-CDP8 peptides are competitive inhibitors (Fig. 3). The results indicate that these peptides interact directly with amino acid residues located in the active site or with amino acids located in the substrate-binding region of the protease, preventing substrate entry to the active site. Since L-CDP1 and L-CDP7 present IC_50_ values < 2 µM, they were considered for profound evaluation.

**Figure 3.**
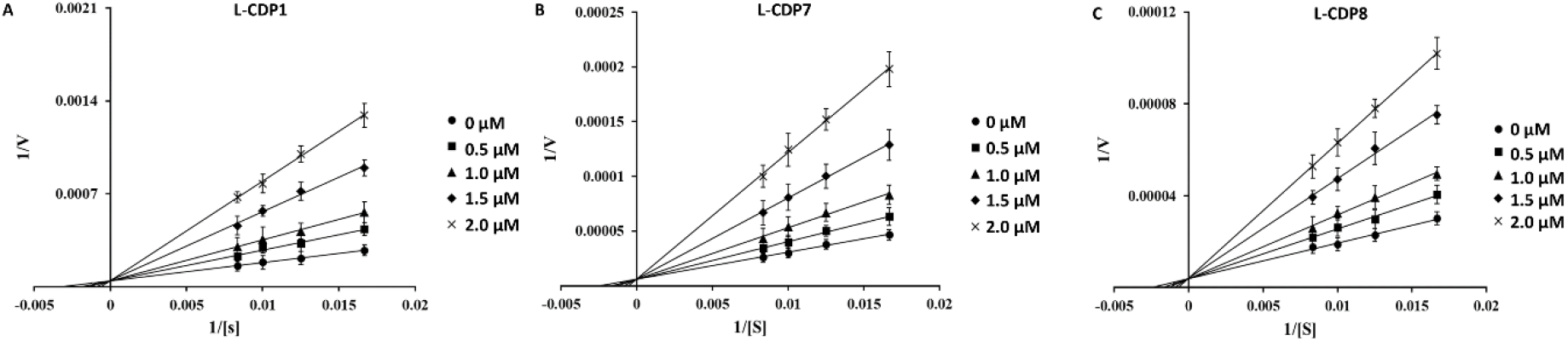
Inhibition mode of L-CDP1, L-CDP7 and L-CDP8 over SARS-CoV-2 3CLpro. Lineweaver-Burk plots to determine the inhibition mode are presented. [S] is the substrate concentration; v is the initial reaction rate. **A:** Lineweaver-Burk plot for L-CDP1 inhibition of SARS-CoV-2 3CLpro. **B:** Lineweaver-Burk plot for L-CDP7 inhibition of SARS-CoV-2 3CLpro. **C:** Lineweaver-Burk plot for L-CDP8 inhibition of SARS-CoV-2 3CLpro. Data shown are the mean ± SD from 3 independent measurements (n=3).

### 2.4. The binding affinity of the L-CDP1 and L-CDP7 using surface plasmon resonance

We further investigated the binding affinity of L-CDP1 and L-CDP7 molecules using Surface Plasmon Resonance (SPR) (Table 2, Supplementary Fig. S7). The protease was passed over a CM-5 sensor chip pre-immobilised with the individual L-peptides (L-CDP1 and L-CDP7). L-CDP1 had an equilibrium rate constant (K_D_) of 65 ± 20.1 nM, whereas L-CDP7 shows 304 ± 70.3 nM, also illustrated in Table 2. From the K_D_ value obtained from SPR, it was clearly observed that 3CL^pro^ had a higher binding efficacy for the L-CDP1 receptor.

**Table 2.**
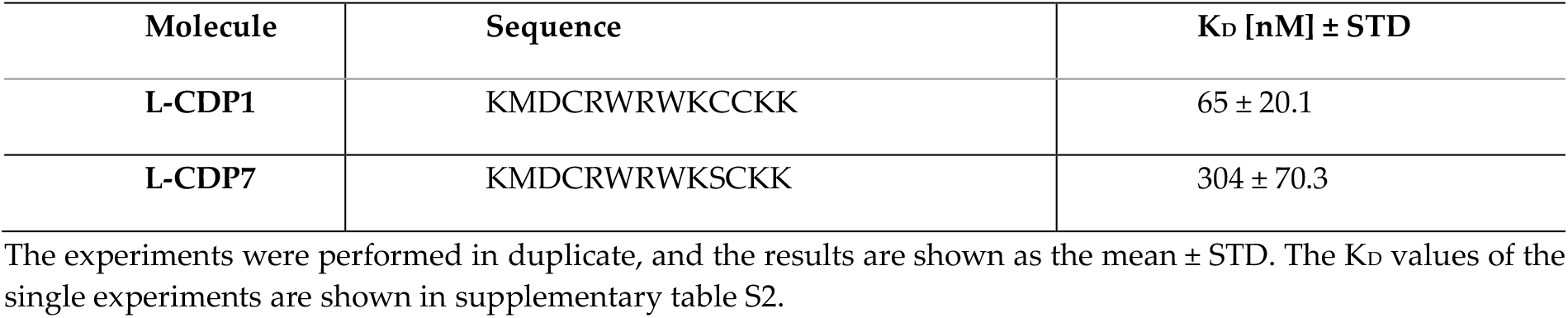
SARS-CoV-2 3CL^pro^ SPR Experiments by L-CDP1 and L-CDP7.

The observed changes in the dissociation process can be attributed to a distinct aspect that the substitution of a sulfur-containing amino acid (Cysteine) by a hydroxyl group (Serine) change the secondary structure (as described before), as well as the mode of interaction of the peptide with the target protease. Jha et al. described in an internalisation study using a Cro derivative peptide (CyLoP-1) that the substitution or deletion of the cysteine residue reduces cellular uptake and cytosolic distribution [30]. Both studies show the importance of the cysteine residue, not only for internalisation but also for inhibiting the SARS-CoV-2 protease.

### 2.5 Inhibition assay of the CDP1 and CDP7 D-peptide enantiomers

In order to conserve all the essential biological properties of the L-enantiomers peptides against human protease degradations, CDP1 and CDP7 were synthesised in D-enantiomers form. D-peptides, when compared to their L-enantiomeric equivalents, possess several therapeutic advantages. As shown previously, the proteolytic stability of D-peptides is superior to L-peptides, which can dramatically increase serum half-life. [42,43], resulting also in reduced immunogenicity and increased bioavailability of D-peptides [44]. Welch et al. 2007 described promising D-peptide inhibitors that effectively inhibit human immunodeficiency virus 1 (HIV-1) entry [45].

L-CDP peptides are susceptible to hydrolysis by proteases, which restrict their therapeutic utility. Based on the mirror symmetry, CDP1 and CDP7 were synthesised (Genscript) in D-enantiomeric form seeking stability. To confirm that the D-peptide were in the correct enantiomeric form, we performed circular dichroism (CD) measurements of both D-CDP1 and D-CDP7 in solution. As expected, the CD spectra of the D-peptides is the mirror image of that of L-peptides (Fig. 4)

**Figure 4.**
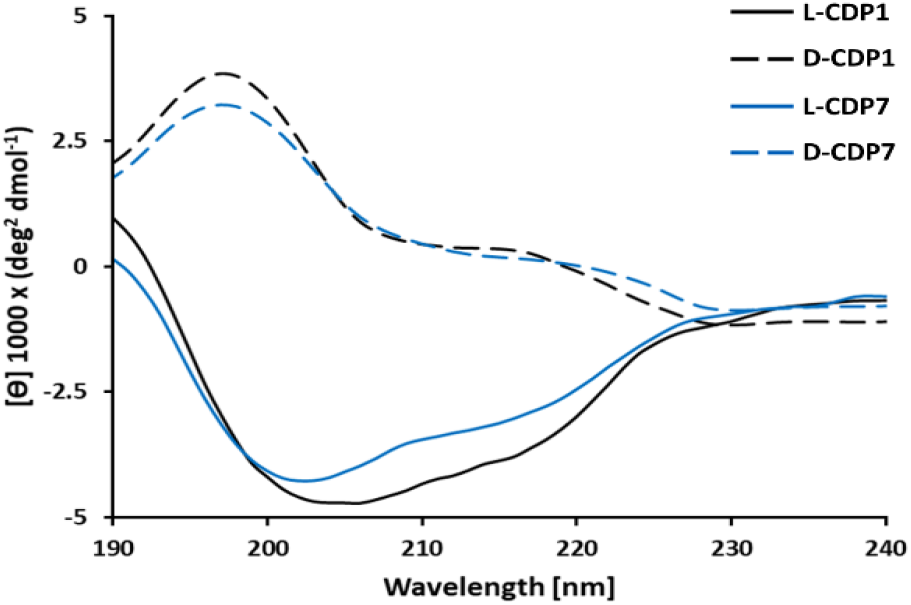
Circular dichroism (CD) spectroscopy of D-CDP peptides. The CD spectrum of D-peptide in solution is presented as molar ellipticity [θ].

The corresponding D-CDP1 and D-CDP7 should specifically bind to the natural L-target with similar affinity. The inhibitory effect, IC_50_, inhibition mode, and K_D_ were determined (Fig. 5 and Supplementary Fig. S8).

**Figure 5.**
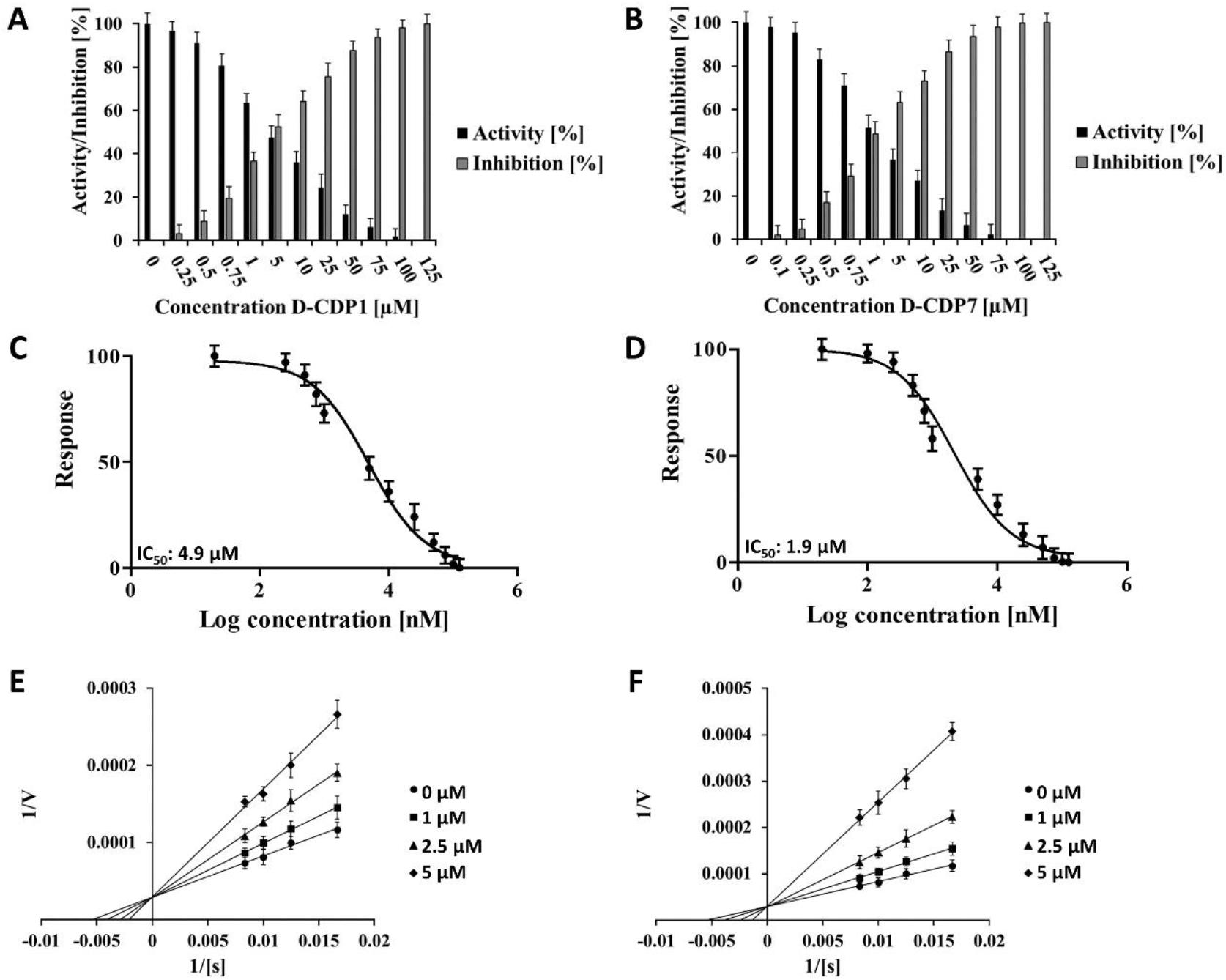
Inhibition effect, dose response curve and inhibition mode of D-CDP1 and D-CDP7 over SARS-CoV-2 3CLpro. **A:** Normalised activity and inhibition of SARS-CoV-2 3CLpro under D-CDP1 influence. **B:** Normalised activity and inhibition of SARS-CoV-2 3CLpro under D-CDP7 influence. **C:** Dose response curve of D-CDP1 and SARS-CoV-2 3CLpro. The normalised response [%] of SARS-CoV-2 3CLpro is plotted against the Log of the D-CDP1 concentration. **D:** Dose response curve of D-CDP7 and SARS-CoV-2 3CLpro. **E:** Lineweaver-Burk plot for D-CDP1 inhibition of SARS-CoV-2 3CLpro. [S] is the substrate concentration; v is the initial reaction rate. **F:** Lineweaver-Burk plot for D-CDP7 inhibition of SARS-CoV-2 3CLpro. Data shown are the mean ± SD from three independent measurements (n=3).

The determined IC_50_ values for D-CDP1 (4.9 ± 1.7 µM) and D-CDP7 (1.9 ± 0.3 µM) are slightly higher than that in L-peptides with an increase of two-fold for D-CDP1 (Table 3). Like the L-peptides, D-CDP1 and D-CDP7 reveal the same as before, a competitive inhibition mode (Fig. 5 E and F).

**Table 3.** Affinity and Inhibitory Activity of the D-CDP1 and D-CDP-7 Peptides.

| Molecule | IC<sub>50</sub> [$\mu\text{M}$] | Inhibition type | K<sub>D</sub> [nM] ± STD |
| --- | --- | --- | --- |
| D-CDP1 | $4.9 \pm 1.7$ | Competitive | $185.7 \pm 17.8$ |
| D-CDP7 | $1.9 \pm 0.3$ | Competitive | $1951.3 \pm 87.5$ |

The MST experiments were performed in triplicates, and the results are shown as the mean ± STD. The K<sub>D</sub> values of the single experiments are shown in supplementary table S3.

SPR experiments of the D-enantiomers with 3CL^pro^ showed no evaluable results. Therefore, the K_D_ values for the D-CDP1 and -D-CDP7 interaction with the 3CL^pro^ was determined using microscale thermophoresis (MST) (Supplementary Fig. S8), which we did not compare with the results for L-CDPs (SPR) as we modified the method to determine the K_D_.

It is well described that a structural conversion or steric incompatibility in D-enantiomers can negatively influence the inhibition and binding behaviour of the D-peptides with the target proteins [46, 47] or the method of choice to determine the K_D_, which was also observed for D-CDP1 and D-CDP7. However, we have shown that both L-enantiomers (CDP1, CDP7) could be significantly modified without altering their function. The D-CDP1 interaction with 3CL^pro^ is around ten times stronger than D-CDP7; this tendency was also observed for the L-enantiomers (Table 3 and Supplementary Table S1).

### 2.6 24 h Stability and promiscuous assays of L/D-CDP1 and L/D-CDP7

The term “promiscuous” inhibitors describe compounds whose inhibition mechanism involves interacting aggregates of many molecules with the target protein. Classified also as “promiscuous” are redox cycling compounds (RCCs) that generate µM concentrations of hydrogen peroxide (H_2_O_2_) in the presence of strong reducing agents, which is presented in the assay buffer in order to maintain the catalytic activity of cysteine proteases, like 3CL [12]. H_2_O_2_ generated by RCCs can indirectly inhibit the catalytic activity of proteins by oxidising accessible cysteine and or tryptophan that are present in CDP1 and CDP7 in L- and D-enantiomers.

To exclude the capacity of both D-peptides to behave as a RCC, we performed a hydrogen peroxide (H_2_O_2_) assay under the influence of TCEP. Our results demonstrated that L/D-CDP1 and L/D-CDP7 do not produce H_2_O_2_ under the influence of 1 mM TCEP and can be excluded as RCCs (Supplementary Fig. S9). Furthermore, a detergent-based control was performed to exclude peptide inhibitors that possibly act as an aggregator of 3CL^pro^, and the experiment was performed by adding 0.001%, 0.01% and 0.1% Triton X-100 to the reaction. Supposed that a molecule would exhibit significant inhibition of 3CL^pro^, which is diminished by detergent, it is almost certainly acting as an aggregation-based inhibitor, as described before [11], which was not observed for L/D-CDP1 and L/D-CDP7 (Supplementary Fig. S10), discarding the possible aggregation properties of the peptides.

Many peptide-based inhibitors lose the inhibitory effect over the time. The stability of the L-and D-peptides (CDP1 and CDP7) selected in this survey was tested over 24 h. The results demonstrated a constant inhibition of SARS-CoV-2 3CL^pro^ over 24 h by both peptides, demonstrating their stability over time and is not prone to digestion by the protease. (Supplementary Fig. S11).

### 2.7 Cytotoxicity assay of D-CDP1 and D-CDP7

A critical factor in evaluating the eligibility of potential lead peptides is cytotoxicity. Cytotoxicity of D-CDP1 and D-CDP7 conducted on the Vero cell line showed > 80% viability at the Minimum Inhibitory Concentration (MIC) concerning the untreated cells (Supplementary Fig. 12). The cells were treated with D-CDP1 and D-CDP7 at concentrations ranging from 0.2 to 98 µM. The results showed that both molecules have no cytotoxic effect at the calculated IC_50_ concentrations described above (Supplementary Fig. S12). The low cytotoxicity of both D-peptides agrees with the results described before for the L-peptides [30].

### 2.8 Antiviral activity of D-CDP1 and D-CDP7

To further substantiate the enzyme inhibition results, we evaluated the ability of these peptides to inhibit SARS-CoV-2. To test whether the two SARS-CoV-2 3CL^pro^ inhibitors, D-CDP1 and D-CDP7, were able to inhibit SARS-CoV-2 replication in cell culture, an African green monkey cell line that supports productive SARS-CoV-2 replication was used. These Vero cells were pre-treated with the two inhibitors for 1 h; subsequently, the cells were infected at an MOI of 0.05. Viral replication was analysed by determining the SARS-CoV-2 RNA within the cell culture supernatant 2 dpi (days post-infection). Pre-treatment at a non-toxic concentration (50 µM) resulted in a significant decrease in viral RNA after 2 dpi (Fig. 6), suggesting that both SARS-CoV-2 3CL^pro^ inhibitors, D-CDP1 and D-CDP7, contribute to impaired SARS-CoV-2 replication through their anti-3CL^pro^ activity.

**Figure 6.**
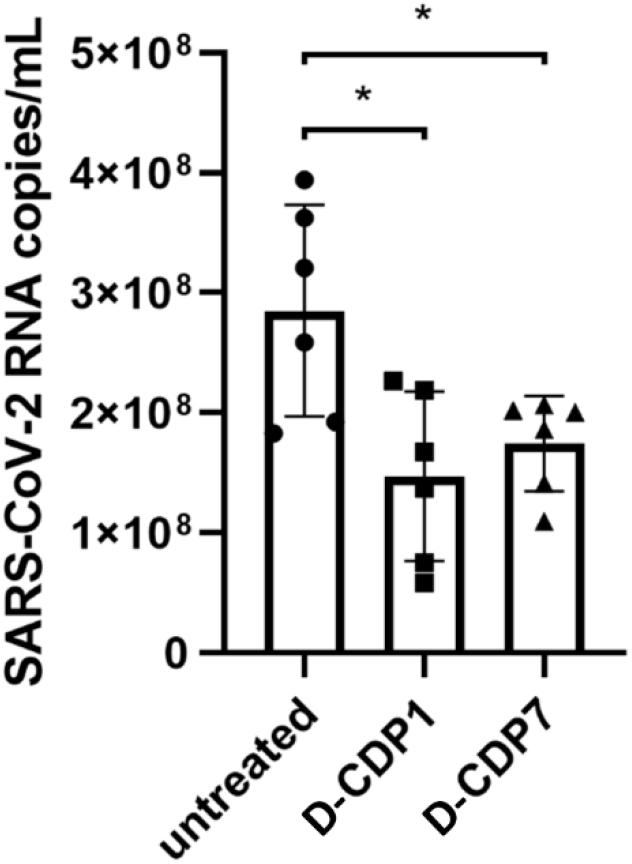
Inhibition of SARS-CoV-2 by D-CDP1 and D-CDP7. Vero cells were treated with 50 µM of either D-CDP1 or D-CDP7, and after an hour of incubation, the cells were infected with SARS-CoV-2 using a multiplicity of infection (MOI) of 0.05. Untreated cells were used as control. Viral RNA in the supernatant was analysed in-house. The graph shows individual data points with mean ± SD (n = 6).

This experiment demonstrates that both D-peptides can impair SARS-CoV-2 replication, most likely by explicitly targeting the viral protease 3CL^pro^. While a twofold reduction in viral RNA, as seen after treatment with D-CDP1, may not seem much regarding the high replication rate, it clearly shows that our compounds exert antiviral activity within SARS-CoV-2-infected cells. These results provide the first proof that the peptides target the inhibition of the SARS-CoV-2 replication. Whether this effect is due to the anti-3CLpro activity and whether the inhibitory effect observed can be optimised by additional modifications is subject to further investigation.

Future work should aim to reveal the mechanism behind the action of both D-peptides and the optimisation of the amino acid composition.

### 2.9 Docking and Molecular Dynamics Simulation

Based on the mode of inhibition studies, protein-peptide (L-, D-CDP1) docking was performed applying the HADDOCK web server program. The atomic coordinates of SARS-CoV-2 3CLpro (PDB entry: 6M2Q) and the atomic coordinates of the peptides were submitted to the platform. The selected clusters were subject to 50 ns of molecular dynamics simulation (MD) in an octahedral TIP3P water box. The flexibility of the protease/peptides structure of the MD simulation system was monitored by calculating the RMSD, RMSF, RoG, and the surface area (Supplementary Fig. S13). The complex analyses of the L-,D-CDP1–3CL complex—through MD simulation—showed constant fluctuation over the 50 ns mostly related to D-CDP1, which is most likely regarding the C-terminus fluctuation (Supplementary Fig. S13B). Based on energy interactions, further analysis of the MD simulations identified the amino acid residues of the protease (below −1 kcal/mol) involved in the binding with the ligand (Fig. 7A).

**Figure 7.**
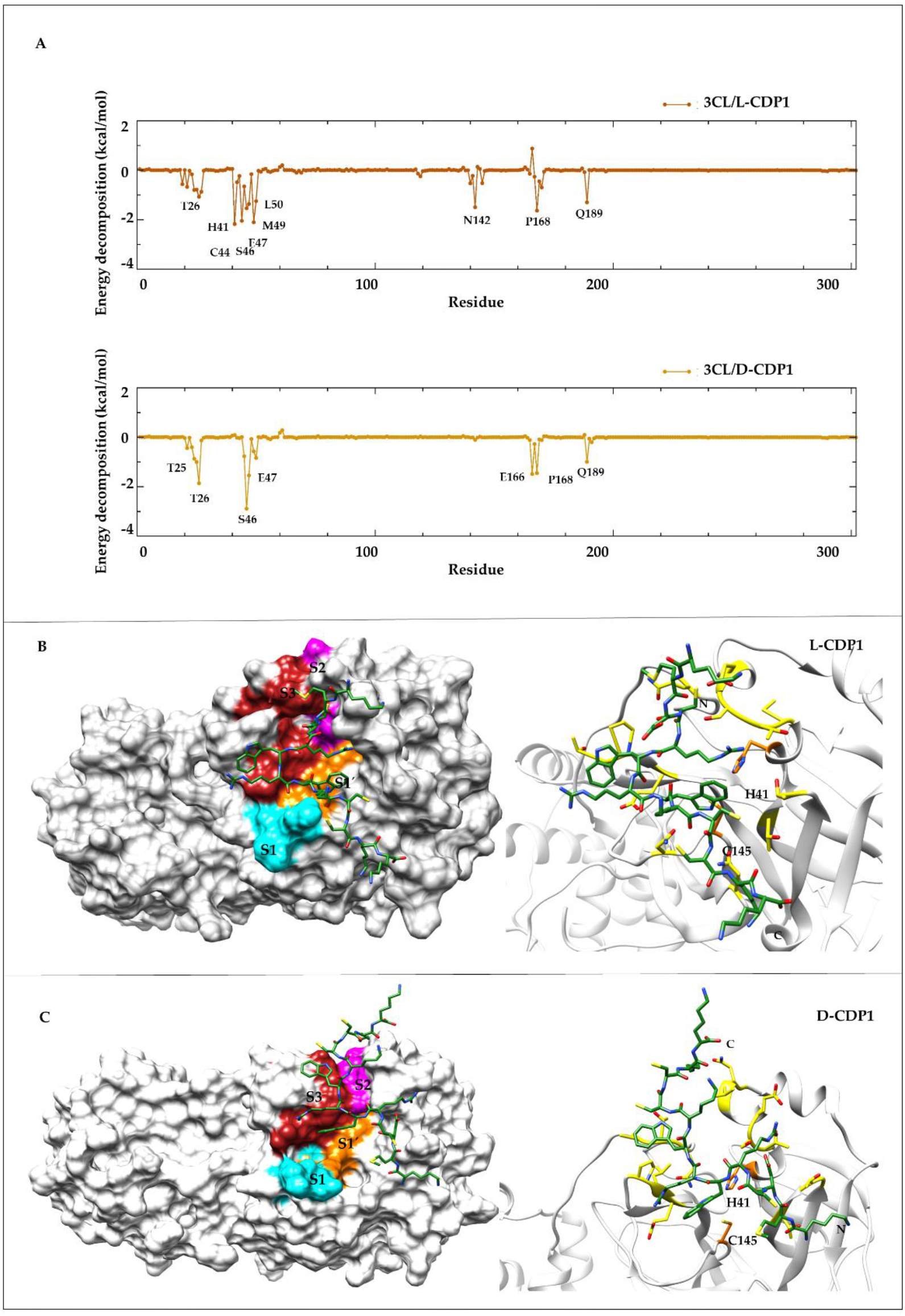
Amino acid residues contributing in the SARS-CoV-2 3CLpro with L-, D-CDP1 peptide interaction. **A)** The energy decomposition of the amino acids from the 3CLpro involved in the interaction with the peptides (upper: L-CDP1, lower: D-CDP1). The results were calculated from MD simulation and was selected with the parameter value below −1 kcal/mol. **B)** Left side, surface representation of the 3CLpro in complex with the L-CDP1 peptide (sticks), the substrate binding sites are colored according: S1 cyan; S1’orange; S2 magenta; S3 red. Right side, cartoon representation of the protease in complex with the peptide (sticks), the amino acids of the catalytic site are shown in sticks (orange). **C**) Left side, surface representation of the 3CLpro in complex with the D-CDP1 peptide (sticks), the substrate binding sites are colored according: S1 cyan; S1’orange; S2 magenta; S3 red. Right side, cartoon representation of the protease in complex with the peptide (sticks), the amino acids of the catalytic site are shown in sticks (orange).

MD simulation revealed a possible mode of interaction of the CDP1 peptides with SARS-CoV-2 3CL^pro^. Therefore, His41, the amino acid residue of the active site interacts through hydrogen bond with one of the residues (Lys31) of the L-CDP1, and the other residues are accommodated in the substrate binding regions (Figure 7B), confirming the competitively mode of interaction, as already described by experimental results (Fig. 3A, 5E and Table 3). However, D-CDP1 does not interact with the amino acids residues of the catalytic dyad; nevertheless, the peptide is seated in the substrate binding region blocking the interaction of the substrate with the active site of the protease (Fig. 7B). Using PDBsum platform the interaction type and amino acid residues involved in the interaction between 3CL^pro^ with the L-or D-CDP1 was identified (Table 4 and Supplementary Fig. S14).

**Table 4.**
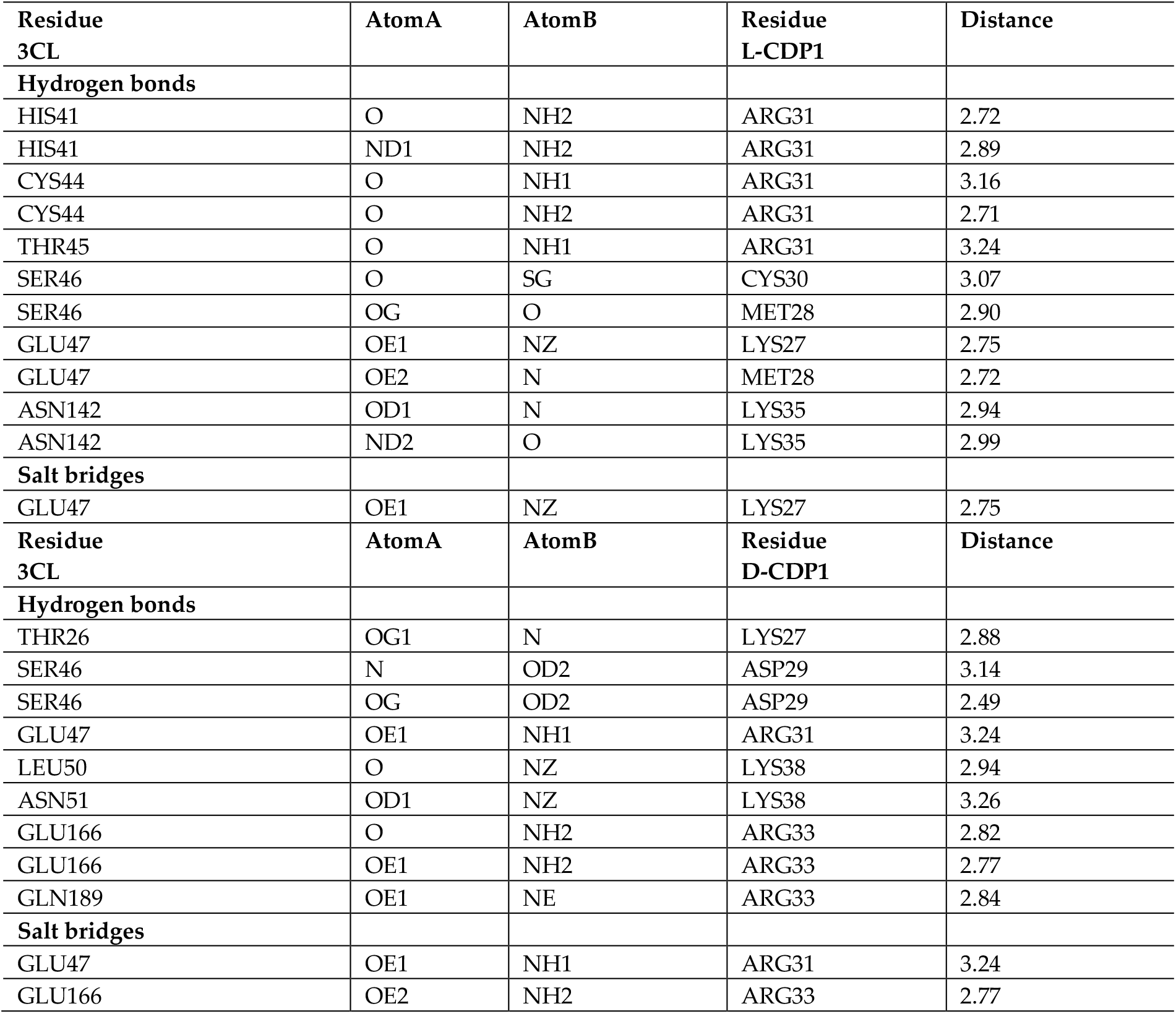
Hydrogen bonds and salt bridges contacts between 3CL^pro^ with L-, D-CDP1 peptides

## 3. Conclusions

Snake venom components have shown great potential for developing lead compounds for new drugs and can be considered mini-drug libraries. For instance, Captopril (Enalapril), Integrilin (Eptifibatide) and Aggrastat (Tirofiban) are FDA approved drugs based on snake venoms. Defibrase/Reptilase (Batroxobin) Hemocoagulase and Exanta (Ximelagatran) are not clinically approved in the US but have been approved for use in other countries. Several venom-derived-drugs are involved in preclinical or clinical trials for a variety of therapeutic applications. In summary, our results demonstrated the promising uses of peptide inhibitors (D-CDP1 and D-CDP7) designed from a polypeptide of snake venom. The advantage of the selected wild type was because of their cell penetration properties, even in D-enantiomer form, high stability and specificity, as well as selectivity against the target 3CL protease was observed. However, in particular, more investigation into the unquestionable untapped therapeutic potential of the selected D-peptide is still required, as well, future optimization.

## 4. Materials and Methods

### 4.1 Crotamine purification

The purification of Crotamine has been described previously [31]. Briefly, Crotamine from crude *Crotalus durissus terrificus* venom obtained from CEVAP (Center for the Study of Venoms and Venomous Animals), Botucatu, Brazil, was isolated by a single cation-exchange chromatography step by utilising a MonoS HR 10/10 column (Amersham Biosciences).

### 4.2 Crotamine derivative peptides (CDP) synthesis

Synthetic Crotamine derivative peptides (CDP) in the L- and D-enantiomeric conformations were synthesised by Genscript (Leiden, NL), with a purity of > 95% (Supplementary Fig. S15). The peptides were acetylated at the N-terminus and methylated at the C-terminus. Essential information about the CDPs used in this study is summarised in Supplementary Fig. S2, S3 and Table S2.

### 4.3 Cloning, expression and purification of SARS-CoV-2 3CL^pro^

SARS-CoV-2 3CL^pro^ (Uniprot entry: P0DTD1, virus strain: hCoV-19/Wuhan/WIV04/2019) was cloned, expressed and purified as described previously [32].

### 4.4 Activity assay of SARS-CoV-2 3CL^pro^

SARS-CoV-2 3CL^pro^ activity assay was performed using a fluorogenic substrate DABCYL-KTSAVLQ↓SGFRKME-EDANS (Bachem, Switzerland) in a buffer containing 20 mM Tris pH 7.2, 200 mM NaCl, 1 mM EDTA, and 1 mM TCEP [32-35]. The reaction mixture was pipetted in a Corning 96-Well plate (Sigma Aldrich) consisting of 0.5 µM protein. The assay was initiated with the addition of the substrate at a final concentration of 50 µM. The fluorescence intensities were measured at 60 s intervals over 30 minutes using an Infinite 200 PRO plate reader (Tecan, Männedorf, Switzerland), the temperature was set to 37 °C. The excitation and emission wavelengths were 360 and 460 nm, respectively.

### 4.5 Inhibition assay of SARS-CoV-2 3CL^pro^

Inhibition of SARS-CoV-2 3CL^pro^ activity by Cro, L- and D-CDPs was investigated using the activity assay described above. 30 µM of the peptides was used for a preliminary screening test.

For the final inhibition assays, 0.5 µM of the protein was incubated with 0-300 µM Cro, 0-150 µM L-CDPs and 0-125 µM D-CDPs, and the mixtures were incubated for 30 minutes at RT. When the substrate with a final concentration of 50 µM was added to the mixture, the fluorescence intensities were measured at 60 s intervals over 30 minutes using an Infinite 200 PRO plate reader (Tecan, Männedorf, Switzerland). The temperature was set to 37 °C, and the excitation and emission wavelengths were 360 and 460 nm, respectively. Inhibition assays were performed in triplicate.

The IC_50_ value was calculated by plotting the initial velocity against various concentrations of the combined molecules using a dose-response curve in GraphPad Prism5 software (Tecan, Männedorf, Switzerland), and data are presented as mean ± SD.

### 4.6 Determination of inhibition mode

The inhibition mode was determined using different final concentrations of the inhibitors and substrate. Briefly, 0.5 µM SARS-CoV-2 3CL^pro^ was incubated with the inhibitor in various concentrations for 30 minutes at RT. Subsequently, the reaction was initiated by the addition of the corresponding concentration series of the substrate. The data were analysed using a Lineweaver-Burk plot; therefore, the reciprocal of velocity (1/V) vs the reciprocal of the substrate concentration (1/[S]) was compared [36,37]. All measurements were performed in triplicate, and data are presented as mean ± SD.

### 4.7 Inhibitor stability over 24 h

Stable inhibition of SARS-CoV-2 3CL^pro^ by L-/D-CDP1 and L-/D-CDP7 was monitored via a 24h inhibition experiment. Briefly, 0.5 µM SARS-CoV-2 3CL^pro^ was incubated with 5 µM of the peptide and incubated for 1/2h, 1h, 2h, 3h, 4h, 5h and 24h at RT. The control was performed with the 3CL^pro^ in the absence of the peptide and measured together after each time point. Subsequently, the reaction was initiated by the addition of the substrate. All measurements were performed in triplicate, and data are presented as mean ± SD.

### 4.8 Assays to exclude L-CDPs and D-CDPs as promiscuous inhibitors

A detergent-based control assay was performed to exclude inhibitors that possibly act as aggregators of the 3CL^pro^ by adding 0.001%, 0.01%, and 0.1% of Triton X-100 to the reaction [38]. Four concentrations of L-CDP1 and D-CDP1 (0.5 µM, 1 µM, 5 µM and 10 µM) and L-CDP7 and D-CDP7 (0.25 µM, 0.5 µM, 1 µM and 5 µM), were tested. All measurements were performed in triplicate, and data are presented as mean ± SD.

Redox cycling compounds (RCCs) generate H_2_O_2_ in the presence of potent reducing agents (e.g. DTT or TCEP). We performed a colourimetric assay to exclude L-CDP1, D-CDP1 and L-CDP7, D-CDP7 as a compound that induces redox cycling in reducing environments.

The assay was performed in Nunc 96-Well plates with a flat bottom (Thermofisher Scientific, USA), and the final volume was 60 µL. 0-60 µM L-CDP1, D-CDP1 and L-CDP7, D-CDP7 were tested in the same activity assay buffer (as described above), 1 mM TCEP was added separately. HRP-PR and 100 µM H_2_O_2_ was used as control. The HRP-PR detection reagent (100 µg/mL phenol red and 60 µg/ml HRP, final) was prepared in Hank’s balanced salt solution (HBSS). HPR-PR and PR without the addition of H_2_O_2_ were used as negative controls. The reducing agents could mediate the oxidation of phenol red (PR) (Sigma Aldrich, USA) based on the H_2_O_2_-dependent horseradish peroxidase (HRP (Sigma Aldrich, USA)) mediated oxidation), which produces a change in its absorbance at 610 nm in alkaline pH [39]. L-CDP1, D-CDP1 and L-CDP7, D-CDP7 were incubated with TCEP at RT for 30 min before adding the HRP-PR detection reagent. After an additional incubation period at RT (10 min), the assay was terminated by adding 10 µL of 1N NaOH to all wells. The absorbance of the phenol red was measured at 610 nm using an Infinite 200 PRO plate reader (Tecan, Männedorf, Switzerland). All measurements were performed in triplicate, and data are presented as mean ± SD.

### 4.9 Determination of dissociation constant using surface plasmon resonance

The dissociation constant (K_D_) of L-enantiomeric peptides CDP1 and CDP7 binding to SARS-CoV-2 3CL^pro^ was determined by Surface Plasmon Resonance Spectroscopy (SPR) using a Biacore 8K instrument (GE Healthcare, Uppsala, Sweden). The peptides were immobilised on two separate channels on a series S CM-5 sensor chip (Cytiva, Uppsala, Sweden) by amine coupling. Both flow cells on each channel were activated by a mixture of 50 mM N-Hydroxysuccinimide (NHS) and 16.1 mM N-ethyl-N’-(dimethylaminopropyl) carbodiimide (EDC) (XanTec, Düsseldorf, Germany) for 7 min. The peptides were diluted to 50 µg/mL in 10 mM sodium acetate pH 5 (Merck, Darmstadt, Germany) and injected overflow cell two of each channel to a final signal of 3400 RU for L-CDP1 and 1300 RU for L-CDP7. After the peptides were immobilised, each channel’s ligand and reference flow cells were quenched by a 7-minute injection of 1 M ethanolamine pH 8.5 (XanTec, Düsseldorf, Germany).

The K_D_ multicycle experiments were determined with PBS containing 0.05 % Tween 20 (AppliChem, Darmstadt, Germany) pH 7.4 as the running buffer. The temperature was set at 25 °C with a flow rate of 30 µL/min. Diluted 3CL^pro^ in the running buffer to the desired concentrations range between 2000 nM - 0.9 nM using 1:2 dilution steps. All samples were injected over the flow cells for 180 s, followed by a dissociation phase of 900 s with a running buffer. A regeneration step was performed to ensure complete dissociation of the protease from the peptides, with 45 s injection of a mixture containing 50 mM oxalic acid, 50 mM phosphoric acid, 50 mM formic acid, and 50 mM malonic acid at pH 5. The reference flow cell and buffer injections (c = 0 nM) were used for double referencing of the sensorgrams. The sensorgrams were fitted by the Steady-State Affinity model implemented in the Biacore Insight Evaluation Software for data evaluation. The experiments were performed in duplicate, and the results are shown as the mean ± STD.

### 4.10 Determination of dissociation constant using microscale thermophoresis (MST)

The binding affinity of the D-enantiomeric peptides CDP1 and CDP7 towards SARS-CoV-2 3CL^pro^ was determined using Microscale thermophoresis (MST). Following the manufacturers’ instructions, peptides were first fluorescently labelled with CF633-NHS (Sigma, Darmstadt, Germany). In short, a 100 µmol/L peptide solution was incubated with 500 µmol/L CF633-NHS solutions in 100 mmol/L NaHCO_3_ pH 8.3 at 25 °C for 1 h with slight agitation. After the reaction was completed, the samples were purified using reversed-phase liquid chromatography on an Agilent 1200 system (Agilent Technologies, Santa Clara, California, USA). The samples were loaded onto a Zorbax SB-300 C-18 column (4.6 * 250 mm, 5 µm) (Agilent Technologies, Santa Clara, California, USA). Mobile phases consisted of A: H_2_O + 0.1% Trifluoroacetic Acid (TFA) (Sigma, Darmstadt, Germany) and B: Acetonitrile (Roth, Karlsruhe, Germany) + 0.1 % TFA. A gradient from 15% B to 45% B in 20 min was applied at a flow rate of 1 mL/min, and CF-633 labelled peptides were eluted after 9.2 min, collected and lyophilised (Christ, Alpha 2-4 LSCbasic, Christ, Osterode am Harz, Germany).

Lyophilized CF-633 labelled peptides were dissolved in 100 µL H_2_O. The absorbance of the sample was measured at 280 and 633 nm using a Shimadzu UV 1800 instrument (Shimadzu, Duisburg, Germany) to calculate the concentration of the peptide.

The MST measurement was performed using a serial dilution of the protease (3CL) (10 µmol/L to 19.53 nmol/L) with a 1:1 dilution step in PBS + 0.05% Tween 20. To each sample, CF633-peptide was added to a final concentration of 100 nmol/L. The measurements were performed on a Nanotemper Monolith NT.115 instrument (NanoTemper Technologies, Munich, Germany). For CF-633 as the fluorescent dye, the red LED was used at 20% power. The MST power was set to 40%, and the temperature was kept constant at 25 °C. Data were evaluated with the Thermophoresis with T-Jump evaluation, with the cold region from −1 s to 0 s and the hot region from 20.67 s to 21.67 s. Data were fitted to the K_D_ Model integrated into the analysis software. The experiments were performed in triplicate, and the results are shown as the mean ± STD.

### 4.11 Circular dichroism (CD) spectroscopy

CD measurements were carried out with a Jasco J-1100 Spectropolarimeter (Jasco, Germany). Far-UV spectra were measured in 190 to 260 nm using a peptide concentration of 30 µM in H_2_O. The secondary structure of L-CDP1, L-CDP2, L-CDP7 and L-CDP8 and D-CDP1 and D-CDP7 was checked. A 1 mm path length cell was used for the measurements; 15 repeat scans were obtained for each sample, and five scans were conducted to establish the respective baselines. The averaged baseline spectrum was subtracted from the averaged sample spectrum. The results are presented as molar ellipticity [θ], according to the equation (1):

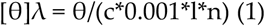

where θ is the ellipticity measured at the wavelength λ (deg), c is the peptide concentration (mol/L), 0.001 is the cell path length (cm), and n is the number of amino acids. Secondary structure prediction based on the CD data was performed with the BeStSeL online tool [40].

### 4.12 Cell viability assay

Cell viability assay was performed using the reduction of [3-(4,5-dimethylthiazol-2-yl)-2,5-diphenyl tetrazolium bromide - MTT] to investigate the cytotoxicity of D-CDP1 and D-CDP7. Therefore, Vero cells were cultivated in DMEM medium supplemented with 2% fetal calf serum (FCS) and 1% penicillin/streptomycin. A total of 2000 cells per well in a volume of 100 µl were seeded on flat-bottomed 96-well plates (VWR, Radnor, USA) and incubated in a 95% humidified atmosphere with 5% CO_2_ at 37 °C for 24 h. Subsequently, the cells were treated with 0.2, 1, 5, 10, 25, 50 and 98 µM of each molecule at 37 °C with 5 % CO_2_ for a period of 24 h.

According to the manufacturer’s instructions, cell viability was measured using the Cell Proliferation Kit I (Roche, Basel, Switzerland) in a 3-fold determination. The absorbance of the formazan product was determined by measuring the absorption at 570 nm subtracted by the absorbance at 690 nm in a microplate reader (BMG Labtech, Ortenberg, Germany). The results were normalised to the mean value of cells treated with medium only. As a negative control for cytotoxicity, 0.1 % Triton X-100 diluted in the medium was used. Cell viability was calculated according to equation (2):

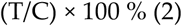

in which T and C represented the optical density of the treated well and control groups, respectively. The MTT assays for D-CDP1 and D-CDP-7 were performed as triplicates, and the results are shown as mean ± SD.

### 4.13 SARS-CoV-2 infection assay

Experiments with SARS-CoV-2 were performed under biosafety level 3 (BSL-3) conditions at the University Hospital Düsseldorf. Vero cells (ATCC-CCL-81, obtained from LGC Standards) were seeded into the wells of a 96-well plate in Dulbecco’s Modified Eagle Medium - DMEM (Gibco, Carlsbad, California) with 2% fetal calf serum (PAN Biotech, Germany) and 1% penicillin/streptomycin (Gibco, Carlsbad, California) at a density of 5×10^3^ cells well^-1^. Following overnight incubation, the medium (100 µl) was replaced with 100 µl new medium containing the respective compounds D-CDP1 or D-CDP7 at a final concentration of 50 µM and the cells were incubated for one hour at 37 °C in a humidified cell culture incubator. Cells were infected with a SARS-CoV-2 B1.1 isolate (SARS-CoV-2 NRW-42 isolate; GISAID accession number: EPI_ISL_425126) (PMID: 32876341 [41]) at a multiplicity of infection (MOI) of 0.05. At two days post-infection (dpi), quantitative RT-PCR was performed. Therefore, 50 µl cell culture supernatant was mixed with 200 µl AVL buffer (Qiagen, Germany) and, following 10 min of incubation at RT. Subsequently, 200 µl of 100% ethanol was added to the mixture. Following the manufacturer’s instructions, RNA was extracted from 200 µl of the inactivated supernatant using the EZ1 Virus Mini Kit (Qiagen, Germany). As described previously, the 60 µl eluate was analysed via in-house qRT-PCR (PMID: 32876341 [41]). In short, a 113 bp sequence of the SARS-CoV-2 Envelope open reading frame was amplified using the real-time TaqMan technique.

### 4.14 Statistical analysis

All data are expressed as the mean ± the standard deviations (SDs). The statistical significance of the mean values’ differences was assessed with one-way analyses of variance (ANOVA), followed by Tukeys’ multiple comparison test. Significant differences were considered at p < 0.05 (*), p < 0.01 (**) and p < 0.001 (***). A standard two-tailed unpaired t-test was used to analyse the inhibitory effect of the D-CDP1 and D-CDP7 on SARS-CoV-2 replication. All statistical analyses were performed with GraphPad Prism software version 8 (San Diego, CA, USA).

### 4.15 System preparation

The main protease (mpro) was retrieved from pdb databank (PDB code: 6M2Q), and the L-CDP1 was created using the crystal structure of the white type crotamie (PDB ID. 4GV5), the D-CDP1 was generated using the tleap in Amber. Docking was performed using HADDOCK webserver [48]. The top four structures were downloaded and viewed by PyMOL. The H++ web server [49] was used to assign the correct lateral chain protonation of the amino acids to pH 7.0 for the simulated systems. After that, the complex (mpro + L or D-peptide) was placed in an octahedral TIP3P water box with water extended 10 Å away from any solute atom and Cl-ions were added to neutralize the systems.

### 4.16 Simulation setup

All the MD simulations were carried out by Amber18 [50]. Atomic interactions were represented by the FF19SB [51] force field. Starting structures were submitted in a two-step minimization process to remove bad contacts. In the first stage, the complex was restricted and minimized during 5000 steepest descending steps, followed by 5000 conjugated gradient steps, with a force constant of 10.0 kcal/mol-Å². A second round of unconstrained energy minimization was performed during 10,000 steps. The system was slowly heated from 0 to 298 K for 0.5 ns under constant atom number, volume, and temperature (NVT) ensemble, with the complex constrained with a constant force of 10 kcal/mol-Å².

Equilibration was divided into six steps of decreasing constant force for the complex atoms from 10 to 0 kcal/mol-Å², and performed under constant atom number, pressure, and temperature (NPT) ensemble. Lastly, the production run was performed for 50 ns in the NVT ensemble without any restriction. In order to allow a 2 fs dynamic time interval the SHAKE restrictions were applied to all bonds with hydrogen atoms. The Particle mesh Ewald (PME) [52] method, with a 8 Å cutoff, was used to calculate the long-range electrostatics interactions. Temperature (298 K) was controlled by Langevin coupling and pressure (1 atm) was controlled by Berendsen barostat.

### 4.17 Molecular Dynamics Analysis and Interaction Energy Calculation

MD results were analyzed using CPPTRAJ [53] tools for the AmberTools19 package [54]. The Root-Mean-Square Deviation (RMSD) was used to investigate the equilibration and convergence of the simulations. Protein and peptide flexibility was accessed by the Root-Mean-Square Fluctuation (RMSF) of the Cα atoms. Structural changes in the protein were evaluated by the radius of gyration (RoG) and solvent accessible surface area.

The molecular mechanics/generalized Born surface area (MM/GBSA) was calculated between protein–peptide complexes using the generalized Born (GB)-Neck2 [55] implicit solvent model (igb = 8), in the steady-state regime of the entire simulation time, stripping the solvent and ions.

## Supplementary Materials

**Figure S1**. Purification of SARS-Cov-2 3CL^pro^. **Figure S2**. Primary structure of L-CDP1 to L-CDP5. **Figure S3**. Primary structure of L-CDP6 to L-CDP9. **Figure S4**. Inhibition effect of Crotamine and L-CDPs over SARS-CoV-2 3CL^pro^. **Figure S5**. Crotamine and L-CDPs with inhibitory activity against SARS-CoV-2 3CL^pro^. **Figure S6**. Circular dichroism (CD) spectroscopy of L-CDP1, L-CDP2, L-CDP7 and CDP8. **Figure S7**. Dissociation constant (KD) determination of L-CDP1 and L-CDP7 binding to SARS-CoV-2 3CL^pro^ using surface plasmon resonance (SPR). **Figure S8**. Dissociation constant (K_D_) determination of D-CDP1 and D-CDP7 binding to SARS-CoV-2 3CL^pro^ using microscale thermophoresis (MST). **Figure S9**. H_2_O_2_ generating capacity of L- and D-CDP1, CDP7 under the influence of 1 mM TCEP. **Figure S10**. Effect of Triton X-100 on the L- and D-CDP1, CDP7 inhibition against SARS-CoV-2 3CL^pro^. **Figure S11**. 24 h inhibition experiment of L- and D-CDP1, CDP7 against SARS-CoV-2 3CL^pro^. **Figure S12**. MTT assay of D-CDP1 and D-CDP7 on Vero CCL-81 cells. **Figure S13**. Time dependent modifications of the 3CL^pro^/L- D-CDP1 complex. **Figure S14**. List of amino acids-amino acids interactions across 3CLS-peptide interface. **Figure S15**. HPLC chromatogram of the L-CDP and D-CDP peptides. **Table S1**. K_D_ determination of L-CDP1 and L-CDP7 using SPR. **Table S2**. Basic information about tested L- and D-CDPs. **Table S3**. K_D_ determination of D-CDP1 and D-CDP7 using MST.

## Author Contributions

**RJE and MAC** Conceptualisation, Methodology, Investigation, Validation, Formal analysis, Writing. **IG, MT** Methodology, Investigation, Validation. **PNO, LM, OA** and **HS** designed the infection assays with **PNO** and **LM** performing and, **OA** and **HS** supervising the experiments. **MAC** and **DSO** Docking and Molecular Dynamic simulation. **DW** and **RKA** Resources, Reviewing. **MAC** Supervising, Reviewing and Editing.

## Data Availability Statement

All data are reported in the text and supplementary materials.

## Acknowledgements

We would like to thank the support of the Institute of Biological Information Processing (IBI-7) Forschungszentrum Jülich, Germany. We are grateful to our Brazilian partners who are constantly being neglected by the federal government.

## Conflicts of Interest

The authors declare no conflict of interest. The funders had no role in the design of the study, in the collection, analyses, or interpretation of data, in the writing of the manuscript, or in the decision to publish the results.

### Abbreviation used

3CL^pro^: C30 Endopeptidase/3C-like protease
SARS-CoV-2: severe acute respiratory syndrome-coronavirus-2
COVID-19: Coronavirus Disease-2019
Cro: Crotamine
L-CDP: crotamine derivative L-enantiomeric peptide
D-CDP: crotamine derivative D-enantiomeric peptide
WHO: World Health Organization
NSP: Nonstructural proteins
AMP: Antimicrobial peptides
ZIKV: Zika virus
DENV: Dengue virus
IAV: Influenza A virus
CPP: Cell-penetrating polypeptide
SPR: Surface plasmon resonance
K_D_: Dissociation constant
HIV-1: Human immunodeficiency virus 1
RCC: Redox cycling compound
TCEP: Tris(2-carboxyethyl)phosphine hydrochloride
MIC: Minimum Inhibitory Concentration
dpi: Days post-infection
RMSD: Root-mean-square deviation
RMSF: Root-mean-square flutuation
RoG: Radius of gyration
MD: Molecular dynamics
CEVAP: Center for the Study of Venoms and Venomous Animals
HRP-PR: Horseradish peroxidase-phenol red
RT: Room temperature
MST: Microscale thermophoresis
PBS: Phosphate buffered saline
MTT: 3-(4,5-dimethylthiazol-2-yl)-2,5-diphenyl tetrazolium bromide
FCS: Fetal calf serum
DMEM: Dulbecco′s Modified Eagle′s Medium
MOI: Multiplicity of infection
NVT: Number, volume, and temperature
NPT: Number, pressure, and temperature.

## References

1. Lu, H.; Stratton, C.W.; Tang, Y.W. Outbreak of pneumonia of unknown etiology in Wuhan, China: The mystery and the miracle. J. Med. Virol. 2020, 92, 401–402.

2. WHO. 2021. World Health Organization, Coronavirus disease 2019 (COVID-19) Dashboard, 05.11.2021.

3. Polack, F.P.; Thomas, S. J.; Kitchin, N.; Absalon, J.; Gurtman, A.; Lockhart, S.; Perez, J.L.; Marc, G.P.; Moreira, E.D.; Zerbini, C.; Bailey, R.; Swanson, K.A.; Gruber, W. C. Safety and efficacy of the BNT162b2 mRNA Covid-19 vaccine. N. Engl. J. Med. 2020, 383, 2603–2615.

4. Baden, L.R.; El Sahly, H.M.; Essink, B.; Kotloff, K.; Frey, S.; Novak, R.; Diemert, D.; Spector, S.A.; Rouphael, N.; Creech, C.B.; McGettigan, J.; Khetan, S.; Zaks, T. Efficacy and safety of the mRNA-1273 SARS-CoV-2 vaccine. N. Engl. J. Med. 2021, 384, 403–416.

5. Voysey, M.; Clemens, S.A.C.; Madhi, S.A.; Weckx, L.Y.; Folegatti, P.M.; Aley, P.K.; Bijker, E. Safety and ef-ficacy of the ChAdOx1 nCoV-19 vaccine (AZD1222) against SARS-CoV-2: an interim analysis of four randomised controlled trials in Brazil, South Africa, and the UK. The Lancet. 2021, 397, 99–111.

6. Livingston, E.H.; Malani, PN; Creech, C.B. The Johnson & Johnson Vaccine for COVID-19. JAMA 2021, 325, 1575–1575.

7. Madsen, L.W. Remdesivir for the Treatment of Covid-19-Final Report. N. Engl. J. Med. 2020, 338, 1813–1826.

8. Hosseinzadeh, M.H.; Shamshirian, A.; Ebrahimzadeh, M.A. Dexamethasone Vs. COVID-19: An Experimental Study in Line with the Preliminary Findings of a Large Trial. Int. J. Clin. Pract. 2020, e13943.

9. The Japanese Association for Infectious Diseases. Treatment of novel coronavirus disease in Japan (first edition). 2020.

10. French Resuscitation Society. Expert recommendations on treating patients during SARS-CoV-2 epidemic (France). 2020.

11. Italian Society of Infectious and Tropical Diseases (SIMIT). Guidelines for the treatment and support management of patients with COVID-19 coronavirus infection (second edition) (Italy). 2020.

12. Rambaut, A.; Holmes, E.C.; O’Toole, Á.; Hill, V.; McCrone, J.T.; Ruis, C.; du Plessis, L.; Pybus, O.G. A dynamic nomenclature proposal for SARS-CoV-2 lineages to assist genomic epidemiology. Nat. Microbiol. 2020, 5, 1403–1407.

13. Ramajayam, R., Tan, K.P., Liang, P.H. Recent development of 3C and 3CL protease inhibitors for anti-coronavirus and anti-picornavirus drug discovery. Biochem. Soc. Trans. 2011, 39, 1371–1375.

14. Ren, Z.; Yan, L.; Zhang, N.; Guo, Y.; Yang, C.; Lou, Z.; Rao, Z. The newly emerged SARS-like coronavirus HCoV-EMC also has an” Achilles’ heel”: current effective inhibitor targeting a 3C-like protease. Protein Cell. 2013, 4, 248–50.

15. Anand, K.; Palm, G.J.; Mesters, J.R.; Siddell, S.G.; Ziebuhr, J.; Hilgenfeld, R. Structure of coronavirus main proteinase reveals combination of a chymotrypsin fold with an extra α-helical domain. EMBO J. 2002, 21, 3213–3224.

16. Anand, K.; Ziebuhr, J.; Wadhwani, P.; Mesters, J.R.; Hilgenfeld, R. Coronavirus main proteinase (3CLpro) structure: basis for design of anti-SARS drugs. Science. 2003, 300, 1763–1767.

17. Yang, H.; Yang, M.; Ding, Y.; Liu, Y.; Lou, Z.; Zhou, Z.; Sun, L.; Mo, L.; Ye, S.; Pang, H.; Gao, G.F.; Anand, K.; Bartlam, M.; Hilgenfeld, R.; Rao, Z. The crystal structures of severe acute respiratory syndrome virus main protease and its complex with an inhibitor. PNAS. 2003, 100, 13190–13195.

18. Pillaiyar, T.; Manickam, M.; Namasivayam, V.; Hayashi, Y.; Jung, S.H. An overview of severe acute respiratory syndrome–coronavirus (SARS-CoV) 3CL protease inhibitors: peptidomimetics and small molecule chemotherapy. J. Med. Chem. 2016, 59, 6595–6628.

19. Ahmed, A.; Siman-Tov, G.; Hall, G.; Bhalla, N.; Narayanan, A. Human antimicrobial peptides as therapeutics for viral infections. Viruses, 2019, 11, 704.

20. Boas, LCPV; Campos, M.L.; Berlanda, R.L.A.; de Carvalho Neves, N.; Franco, O.L. Antiviral peptides as promising therapeutic drugs. Cell Mol. Life Sci. 2019, 76, 3525–3542.

21. Hsieh, I.N.; Hartshorn, K.L. 2016. The role of antimicrobial peptides in influenza virus infection and their potential as antiviral and immunomodulatory therapy. Pharmaceuticals, 2016, 9, 53.

22. Gonçalves, J.M.; Polson, A. The electrophoretic analysis of snake venom. Arch. Biochem. 1947, 13, 253–259.

23. Gonçalves, J.M.; Vieira, L.G. Estudos sobre venenos de serpentes brasileiras I. Análise eletroforética. An. Acad. Bras. Cienc. 1950, 22, 141–150.

24. Hayashi, M.A.; Oliveira, E.B.; Kerkis, I.; Karpel, R.L. Crotamine: a novel cell-penetrating polypeptide nanocarrier with potential anti-cancer and biotechnological applications. Methods Mol. Biol. 2012, 906, 337–352.

25. Yamane, E.S.; Bizerra, F.C.; Oliveira, E.B.; Moreira, J.T.; Rajabi, M.; Nunes, G.L.; de Souza, A.O.; da Silva, I.D.; Yamane, T.; Karpel, R.L.; Silva Jr., P.I.; Hayashi, M.A. Unraveling the antifungal activity of a South American rattlesnake toxin crotamine. Biochimie 2013, 95, 231–240.

26. Nascimento, F.D.; Hayashi, M.A.; Kerkis, A.; Oliveira, V.; Oliveira, E.B.; Rádis-Baptista, G.; Nader, H.B.; Yamane, T.; Tersariol, I.L.S.; Kerkis, I. Crotamine mediates gene delivery into cells through the binding to heparan sulfate proteoglycans. J. Biol. Chem. 2007, 282, 21349–21360.

27. Hayashi, M.A.; Nascimento, F.D.; Kerkis, A.; Oliveira, V.; Oliveira, E.B.; Pereira, A.; Rádis-Baptista, G.; Nader, H.B.; Yamane, T.; Kerkis, I.; Tersariol, I.L. Cytotoxic effects of crotamine are mediated through lysosomal membrane permeabilisation. Toxicon 2008, 52, 508–517.

28. Chen, P.C.; Hayashi, M.A.; Oliveira, E.B.; Karpel, R.L. DNA-interactive properties of crotamine, a cell-penetrating polypeptide and a potential drug carrier. PLoS One 2012, 7, 48913.

29. Kerkis, A.; Kerkis, I.; Radis-Baptista, G.; Oliveira, E.B.; Vianna-Morgante, A.M.; Pereira, L.V.; Yamane, T. Crotamine is a ‘novel cell-penetrating protein from the venom of rattlesnake Crotalus durissus terrificus, FASEB J. 2004, 18, 1407–1409.

30. Jha, D.; Mishra, R.; Gottschalk, S.; Wiesmüller, K.H.; Ugurbil, K.; Maier, M.E.; Engelmann, J. CyLoP-1: a novel cysteine-rich cell-penetrating peptide for cytosolic delivery of cargoes. Bioconjug. Chem. 2011, 22, 319–328.

31. Coronado, M.A.; Georgieva, D.; Buck, F.; Gabdoulkhakov, A.H.; Ullah, A.; Spencer, P.J.; Arni, RK; Betzel, C. Purification, crystallisation and preliminary X-ray diffraction analysis of crotamine, a myotoxic polypeptide from the Brazilian snake Crotalus durissus terrificus. Acta Crystallogr. Sect. F Struct. Biol. Cryst. Commun. 2012, 68, 1052–1054.

32. Eberle, R.J.; Olivier, D.S.; Amaral, M.S.; Gering, I.; Willbold, D.; Arni, RK; Coronado, M.A. The Repurposed Drugs Suramin and Quinacrine Cooperatively Inhibit SARS-CoV-2 3CLpro In Vitro. Viruses. 2021, 13, 873.

33. Zhang, L.; Lin, D.; Sun, X.; Curth, U.; Drosten, C.; Sauerhering, L.; Becker, S.; Rox, K.; Hilgenfeld, R. Crystal structure of SARS-CoV-2 main protease provides a basis for design of improved α-ketoamide inhibitors. Science. 2020, 368, 409–412.

34. Zhang, L.; Lin, D.; Kusov, Y.; Nian, Y.; Ma, Q.; Wang, J.; De Wilde, A. α-Ketoamides as broad-spectrum inhibitors of coronavirus and enterovirus replication: Structure-based design, synthesis, and activity assessment. J. Med. Chem. 2020, 63, 4562–4578.

35. Ma, C.; Sacco, M.D.; Hurst, B.; Townsend, J.A.; Hu, Y.; Szeto, T.; Zhang, X.; Tarbet, B.; Marty, M.T.; Chen, Y.; Wang, J. Boceprevir, GC-376, and calpain inhibitors II, XII inhibit SARS-CoV-2 viral replication by targeting the viral main protease. Cell Res. 2020, 30, 678–692.

36. Roy, A.; Lim, L.; Srivastava, S.; Lu, Y.; Song, J. Solution conformations of Zika NS2B-NS3pro and its inhibition by natural products from edible plants. PLoS One. 2017, 12, e0180632.

37. Motulsky, H.; Christopoulos, A. Fitting models to biological data using linear and nonlinear regression: a practical guide to curve fitting. Oxford University Press. 2004.

38. Feng, B.Y.; Shoichet, B.K. A detergent-based assay for the detection of promiscuous inhibitors. Nat. Protoc. 2006, 1, 550–553.

39. Johnston, P.A. Redox cycling compounds generate H2O2 in HTS buffers containing strong reducing reagents real hits or promiscuous artifactsã Curr. Opin. Chem. Biol. 2011, 15, 174–182.

40. Micsonai, A., Wien, F.; Bulyáki, E.; Kun, J.; Moussong, E.; Lee, Y.H.; Goto, Y.; Réfrégiers, M.; Kardos, J. BeStSel: a web server for accurate protein secondary structure prediction and fold recognition from the circular dichroism spectra. Nucl. Acids Res. 2018, 46, W315–W322.

41. Ramani, A., Müller, L., Ostermann, P.N., Gabriel, E., Abida-Islam, P., Müller-Schiffmann, A., Mariappan, A., Goureau, O., Gruell, H., Walker, A., Andrée, M., Hauka, S., Houwaart, T., Dilthey, A., Wohlgemuth, K., Omran, H., Klein, F., Wieczorek, D., Adams, O., Timm, J., Korth, C., Schaal, H., Gopalakrishnan, J. SARS-CoV-2 targets neurons of 3D human brain organoids. EMBO J. 2020 Oct 15; 39(20):e106230.

42. Van Regenmortel, M.H.; Muller, S. D-peptides as immunogens and diagnostic reagents. Curr. Opin. Biotechnol. 1998, 9, 377–382.

43. Sadowski, M.; Pankiewicz, J.; Scholtzova, H.; Ripellino, J.A.; Li, Y.; Schmidt, S.D.; Mathews, P.M.; Fryer, J.D.; Holtzman, D.M.; Sigurdsson, EM; Wisniewski, T. A synthetic peptide blocking the apolipoprotein E/beta-amyloid binding mitigates beta amyloidtoxicity and fibril formation in vitro and reduces beta-amyloid plaques in transgenic mice. Am. J. Pathol. 2004, 165, 937–948.

44. Dintzis, H.M.; Symer, D.E.; Dintzis, R.Z.; Zawadzke, L.E.; Berg, J.M. A comparison of the immunogenicity of a pair of enantiomeric proteins. Proteins Struct. Funct. Bioinform. 1993, 16, 306–308.

45. Welch, B.D.; VanDemark, A.P.; Heroux, A.; Hill, CP; Kay, M.S. Potent D-peptide inhibitors of HIV-1 entry. Proc. Natl. Acad. Sci. USA. 2007, 104, 16828–16833.

46. Wei, G.; de Leeuw, E.; Pazgier, M.; Yuan, W.; Zou, G.; Wang, J.; Ericksen, B.; Lu, W.Y.; Lehrer, R.I.; Lu, W. Through the looking glass, mechanistic insights from enantiomeric human defensins. J. Biol. Chem. 2009, 284, 29180–29192.

47. Bai, L.; Sheeley, S.; Sweedler, J.V.; Analysis of endogenous D-amino acid-containing peptides in metazoa. Bioanal. Rev. 2009, 1, 7–24.

48. Dominguez, C., Boelens, R., and Bonvin, A. M. J. J. HADDOCK: A Protein™Protein Docking Approach Based on Biochemical or Biophysical Information. J. Am. Chem. Soc. 2003, 125, 1731–1737.

49. Gordon, J.C.; Myers, J.B.; Folta, T.; Shoja, V.; Heath, L.S.; Onufriev A. H++: a server for estimating pKas and adding missing hydrogens to macromolecules. Nucleic Acids Res 2005, 33 (Web Server):W368–71.

50. Case, D.A.; Cerutti, D.S.; Cheatham III, T.E.; Darden, T.A.; Duke, R.E.; Giese, T.J.; Gohlke, H.; Goetz, A.W.; Greene, D.; Homeyer, N.; Izadi, S.; Kovalenko, A.; Lee, T.S.; LeGrand, S.; Li, P.; Lin, C.; Liu, J.; Luchko, T.; Luo, R.; Mermelstein, D.J.; Merz, K.M.; Miau, Y.; Monard, G.; Nguyen, H.; Omelyan, I.; Onufriev, A.; Pan, F.; Qi, R.; Roe, A.; Roitberg, C. et al. D.M. York and P.A. Kollman. 2018, AMBER 2018, University of California, San Francisco.

51. Tian. C.; Kasavajhala, K.; Belfon, K.A.A.; Raguette L.; Huang, H.; Migues, A.N.; et al. ff19SB: Amino-Acid-Specific Protein Backbone Parameters Trained against Quantum Mechanics Energy Surfaces in Solution. J Chem Theory Comput 2020, 16(1):528–52.

52. Darden, T.; York, D.; Pedersen, L. Particle mesh Ewald: An N. log (N) method for Ewald sums in large systems. J Chem Phys 1993, 98(April):10089.

53. Roe, D.R.; Cheatham III, T.E. PTRAJ and CPPTRAJ: software for processing and analysis of molecular synamics trajectory data. J Chem Theory Com 2013, 9(7):3084–95.

54. Case, D.A.; Cheatham, T.E.; Darden, T.; Gohlke, H.; Luo, R.; Merz, K.M. et al. The Amber biomolecular simulation programs. J Comput Chem 2005, 26(16):1668–88.

55. Nguyen, H.; Roe, DR. Simmerling C. Improved Generalized Born Solvent Model Parameters for Protein Simulations. J Chem Theory Comput 2013, 9(4):2020–34.

