## Supplementary Material for "Design of D-amino acids SARS-CoV-2 Main protease inhibitors using the cationic peptide from rattlesnake venom as a scaffold"

**Table of contents**

**Figure S1**. Purification of SARS-Cov-2 3CL$^{pro}$.
**Figure S2**. Primary structure of L-CDP1 to L-CDP5.
**Figure S3**. Primary structure of L-CDP6 to L-CDP9.
**Figure S4**. Inhibition effect of Crotamine and L-CDPs over SARS-CoV-2 3CL$^{pro}$.
**Figure S5**. Crotamine and L-CDPs with inhibitory activity against SARS-CoV-2 3CL$^{pro}$.
**Figure S6.** Circular dichroism (CD) spectroscopy of L-CDP1, L-CDP2, L-CDP7 and CDP8.
**Figure S7**. Dissociation constant (KD) determination of L-CDP1 and L-CDP7 binding to SARS-CoV-2 3CL$^{pro}$ using surface plasmon resonance (SPR).
**Figure S8**. Dissociation constant ($K_D$) determination of D-CDP1 and D-CDP7 binding to SARS-CoV-2 3CL$^{pro}$ using microscale thermophoresis (MST).
**Figure S9**. $H_2O_2$ generating capacity of L- and D-CDP1, CDP7 under the influence of 1 mM TCEP.
**Figure S10**. Effect of Triton X-100 on the L- and D-CDP1, CDP7 inhibition against SARS-CoV-2 3CL$^{pro}$.
**Figure S11**. 24 h inhibition experiment of L- and D-CDP1, CDP7 against SARS-CoV-2 3CL$^{pro}$.
**Figure S12.** MTT assay of D-CDP1 and D-CDP7 on Vero CCL-81 cells.
**Figure S13.** Time dependent modifications of the 3CL$^{pro}$/L- D-CDP1 complex.
**Figure S14.** List of amino acids- amino acids interactions across 3CLS-peptide interface.
**Figure S15.** HPLC chromatogram of the L-CDP and D-CDP peptides.
**Table S1**. $K_D$ determination of L-CDP1 and L-CDP7 using SPR.
**Table S2.** Basic information about tested L- and D-CDPs.
**Table S3**. $K_D$ determination of D-CDP1 and D-CDP7 using MST.

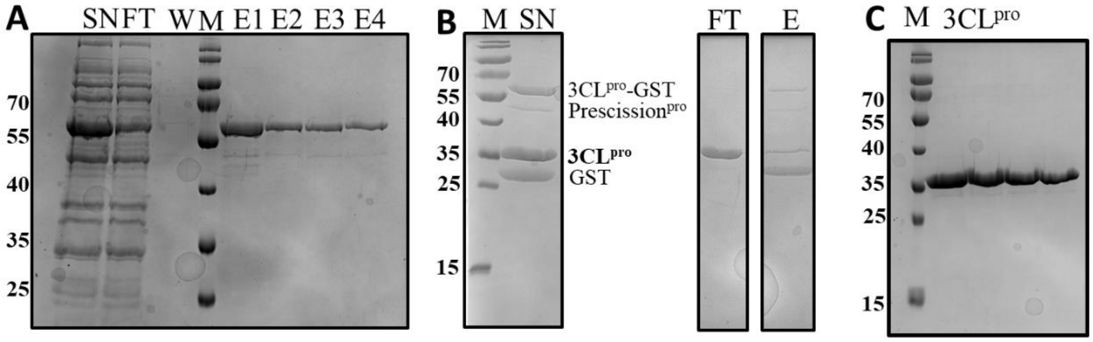


**Figure S1**. Purification of SARS-Cov-2 3CLpro. A: SDS 15% Gel of 3CLpro_GST purification with GSH sepharose. SN: supernatant, FT: flow-through, W: washing step, M: protein marker, E1-E4: Elution. B: SDS 15% Gel of the 3CLpro_GST fusion protein cleavage by PreScissionpro. M: protein marker. SN: cleaving approach, containing 3CLpro_GST, PreScissionpro, 3CLpro and GST. FT: contain 3CLpro, E: elution step including uncleaved 3CLpro_GST, PreScissionpro and GST tag. C: SDS 15% Gel of pure 3CLpro.

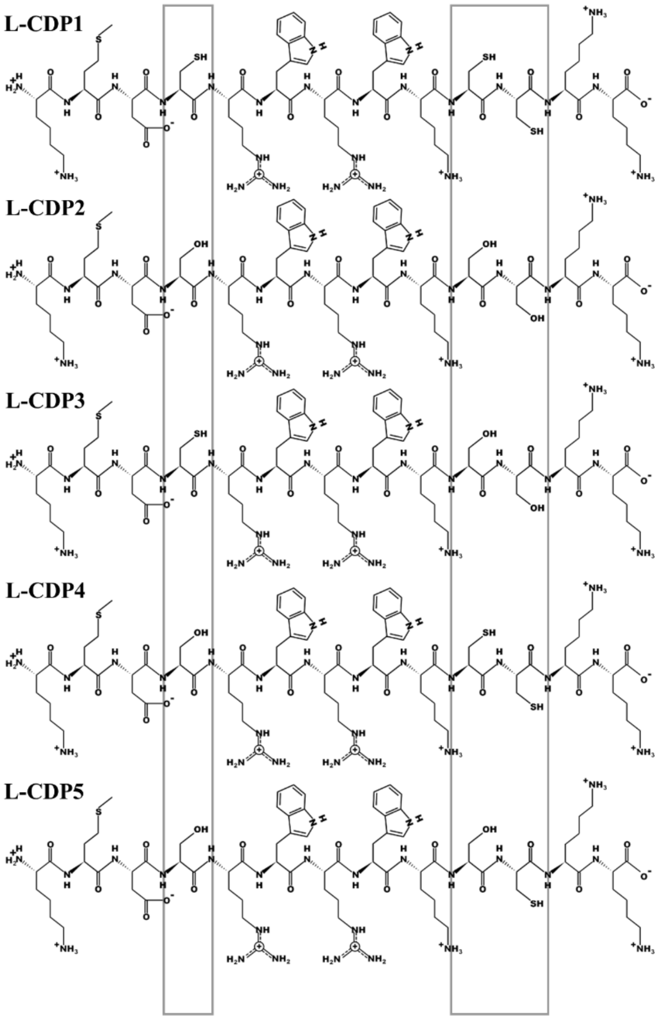


**Figure S2.** Primary structure of L-CDP1 to L-CDP5. Grey boxes label the amino acid variation in the sequences.

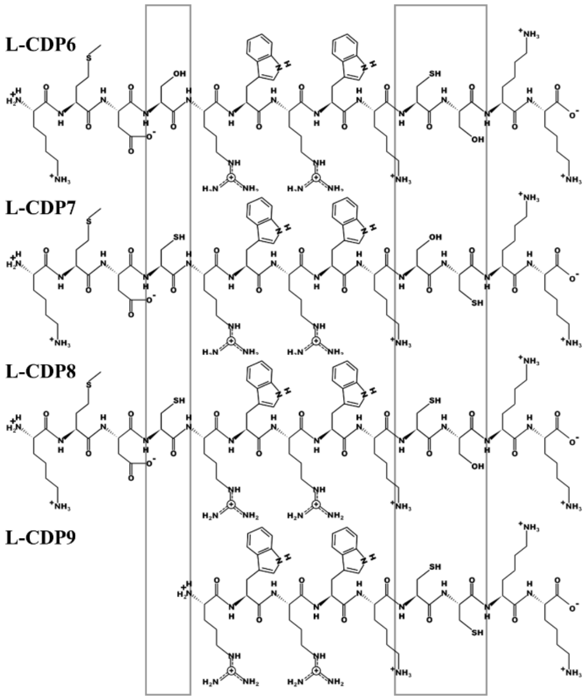


**Figure S3.** Primary structure of L-CDP6 to L-CDP9. Grey boxes label the amino acid variation in the sequences.

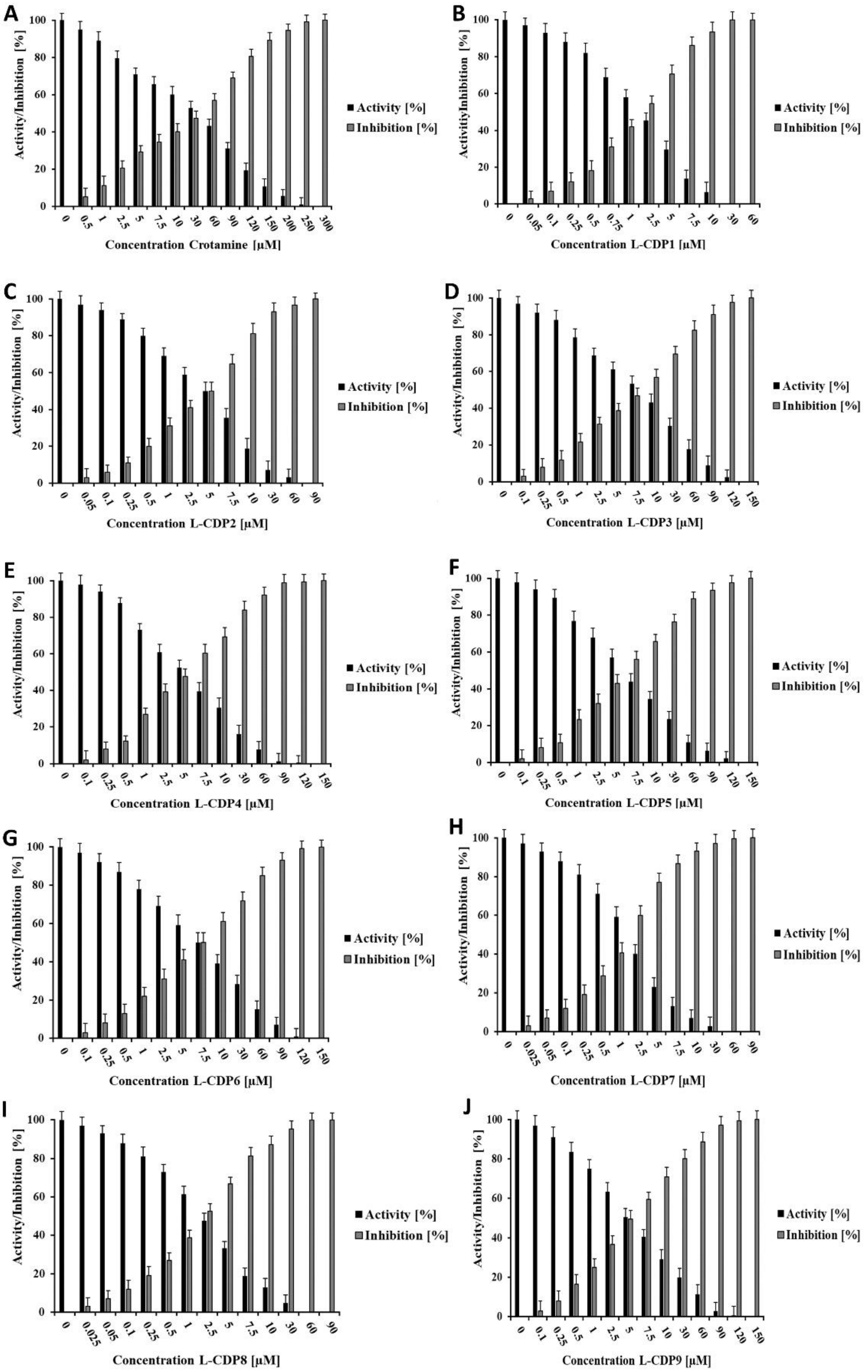


**Figure S4.** Inhibition effect of Crotamine and L-CDPs over SARS-CoV-2 3CL$^{pro}$. Normalized activity and inhibition of SARS-CoV-2 3CL$^{pro}$ under inhibitor influence. **A:** Crotamine, **B:** L-CDP3, **C:** L-CDP4, **D:** L-CDP5, **E:** L-CDP6 and **F:** L-CDP9. Data shown are the mean ± SD from three independent measurements (n=3).

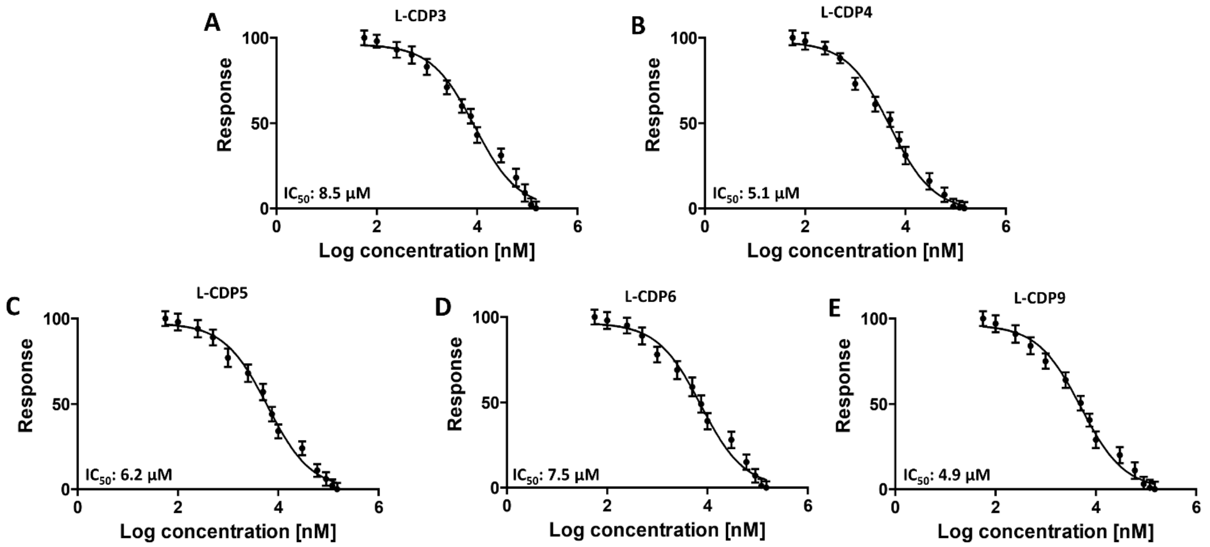


**Figure S5.** Crotamine and L-CDPs with inhibitory activity against SARS-CoV-2 3CL$^{pro}$. Dose-response curves for $IC_{50}$ determination. The normalized response [%] of SARS-CoV-2 3CL$^{pro}$ is plotted against the Log of the inhibitor concentration **A:** L-CDP3, **B:** L-CDP4, **C:** L-CDP5, **D:** L-CDP6, **E:** L-CDP9. Data shown are the mean ± SD from three independent measurements (n=3).

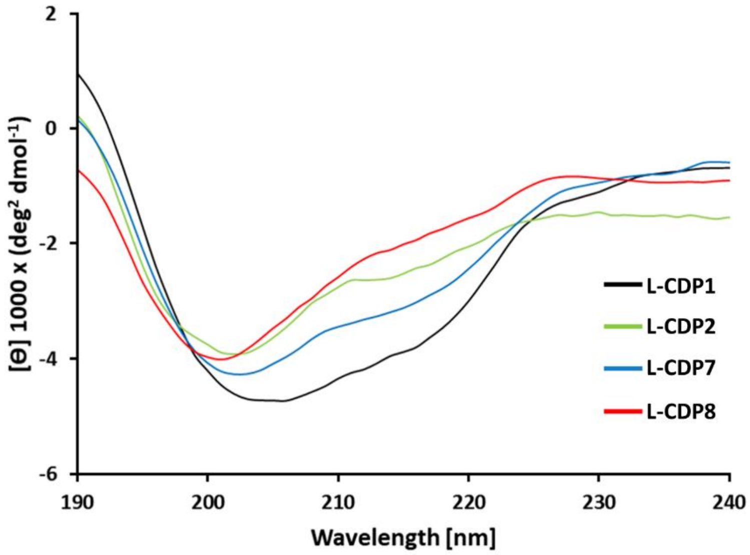


**Figure S6.** Circular dichroism (CD) spectroscopy of L-CDP1, L-CDP2, L-CDP7 and L-CDP8. The CD spectrum of each L-peptide in solution is presented as molar ellipticity [θ].

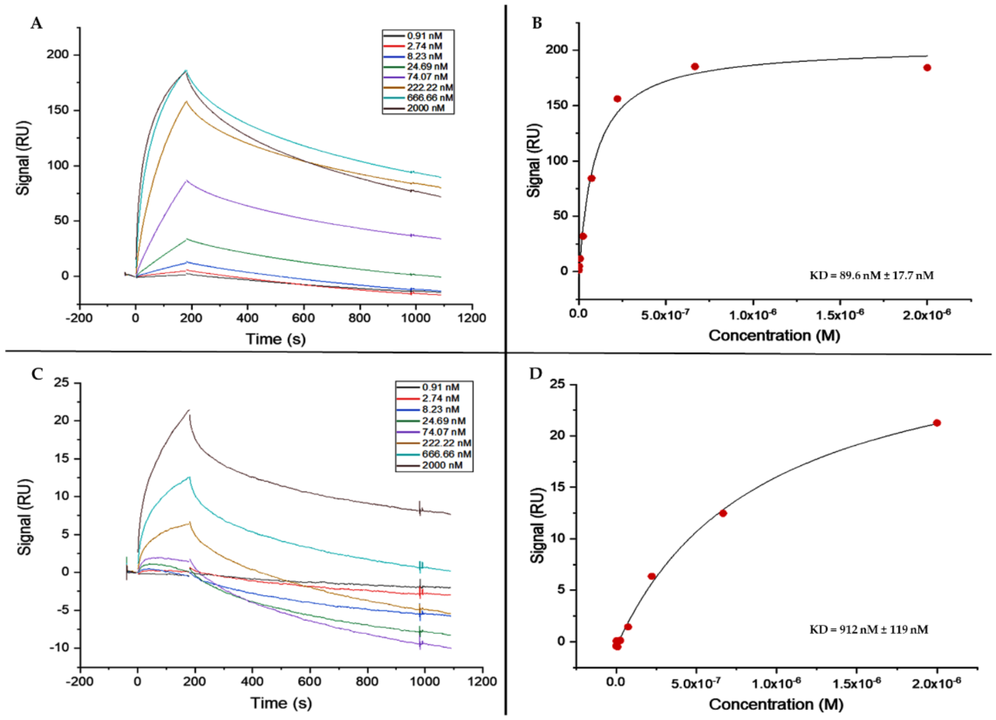


**Figure S7.** Dissociation constant ($K_D$) determination of L-CDP1 and L-CDP7 binding to SARS-CoV-2 $3CL^{pro}$ using surface plasmon resonance (SPR). **A:** SPR sensorgram of $3CL^{pro}$ and L-CDP1. **B:** Saturation curve for the L-CDP1 and $3CL^{pro}$ interaction. **C:** SPR sensorgram of $3CL^{pro}$ and L-CDP7. **D:** Saturation curve for the L-CDP7 and $3CL^{pro}$ interaction.

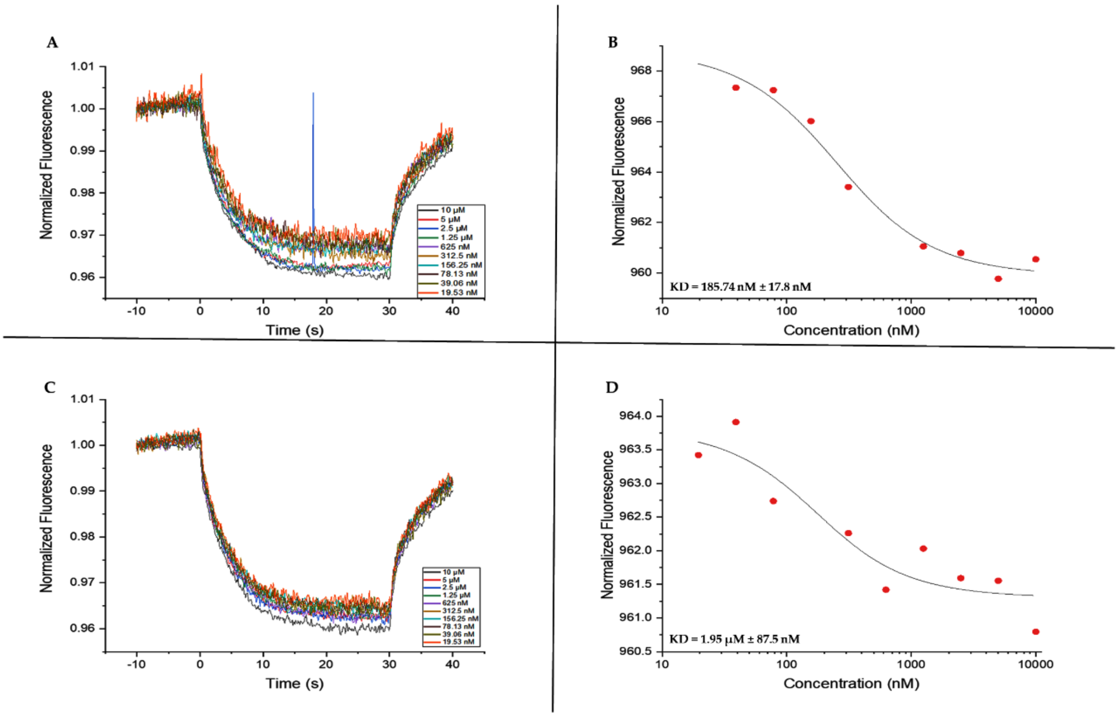


**Figure S8.** Dissociation constant ($K_D$) determination of D-CDP1 and D-CDP7 binding to SARS-CoV-2 $3CL^{pro}$ using microscale thermophoresis (MST). **A:** Thermophoresis data from serial dilutions of D-CDP1. **B:** Binding curve of D-CDP1 with SARS-CoV-2 $3CL^{pro}$. **C:** Thermophoresis data from serial dilutions of D-CDP7. **D:** Binding curve of D-CDP7 with SARS-CoV-2 $3CL^{pro}$.

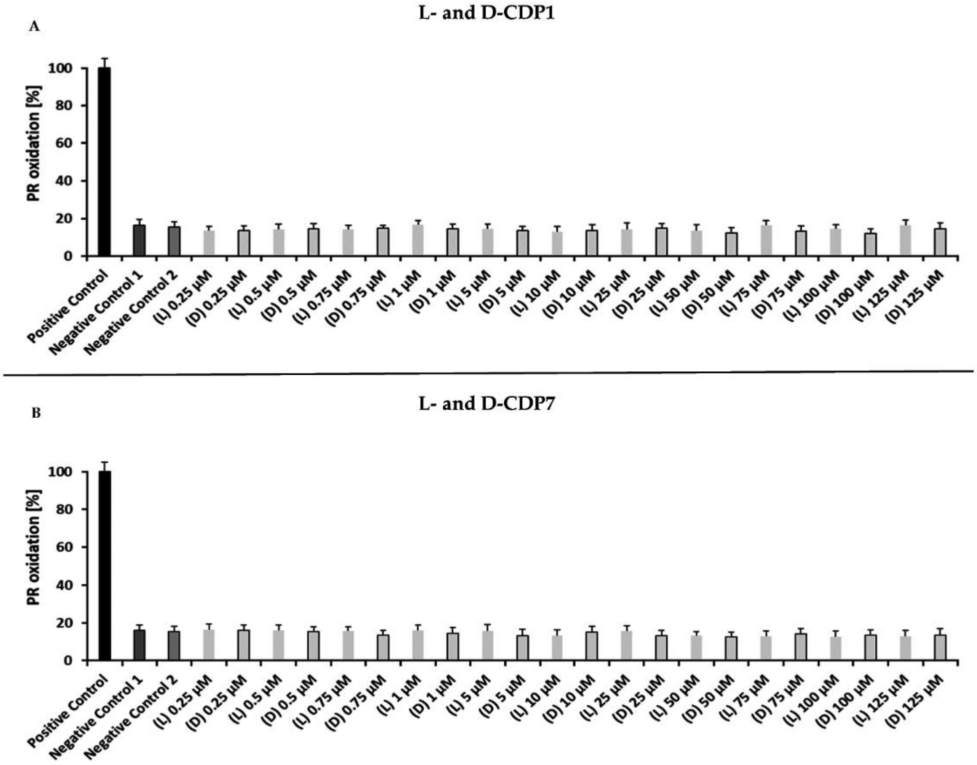


**Figure S9. $H_2O_2$ generating capacity of L- and D-CDPs under the influence of 1 mM TCEP.** The experiments were performed to measure the $H_2O_2$ generating capacity of L-/D-CDP1 and L-/D-CDP7 under the influence of 1mM TCEP. Further, the $H_2O_2$-dependent horseradish peroxidase (HRP) mediates phenol red (PR) oxidation, which can be followed at 610 nm. Positive control: HRP-PR and $H_2O_2$; negative control 1: HRP-PR; negative control 2: PR; Data shown are the mean ± SD from three independent measurements (n=3). **A:** $H_2O_2$ generation by L- and D-CDP1. **B:** $H_2O_2$ generation by L- and D-CDP7.

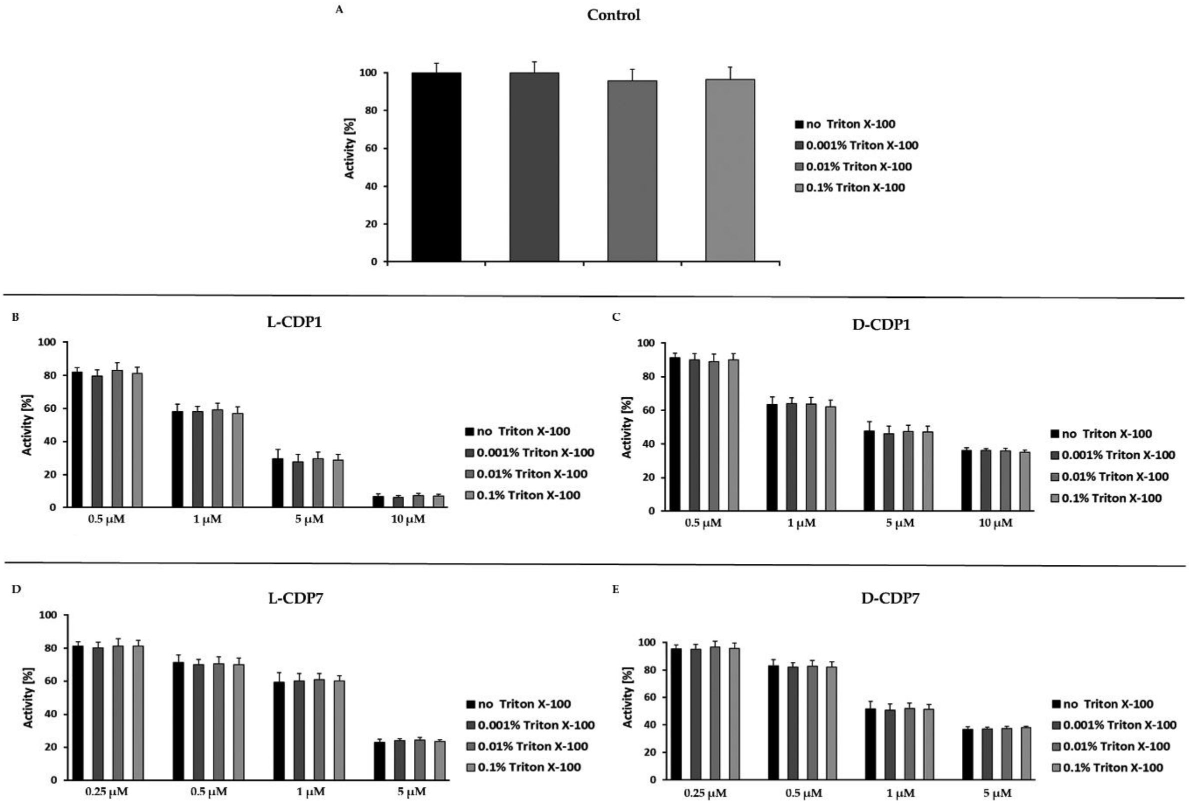


**Figure S10. Effect of Triton X-100 on the L- and D-CDPs inhibition against SARS-CoV-2 3CLpro.** The experiments were performed to exclude the possibility that the peptides inhibit the protease promiscuously by aggregation. Three Triton X-100 concentrations (0.001%, 0.01% and 0.1%) were tested with four different inhibitor concentrations and compared with the results without detergent. Additionally, the effect of Triton X-100 against the protease was tested. Data shown are the mean ± SD from three independent measurements (n=3). **A:** Effect of Triton X-100 on 3CLpro activity. **B:** Effect of Triton X-100 on L-CDP1 inhibition against 3CLpro. **C:** Effect of Triton X-100 on D-CDP1 inhibition against 3CLpro. **D:** Effect of Triton X-100 on L-CDP7 inhibition against 3CLpro. **E:** Effect of Triton X-100 on D-CDP7 inhibition against 3CLpro.

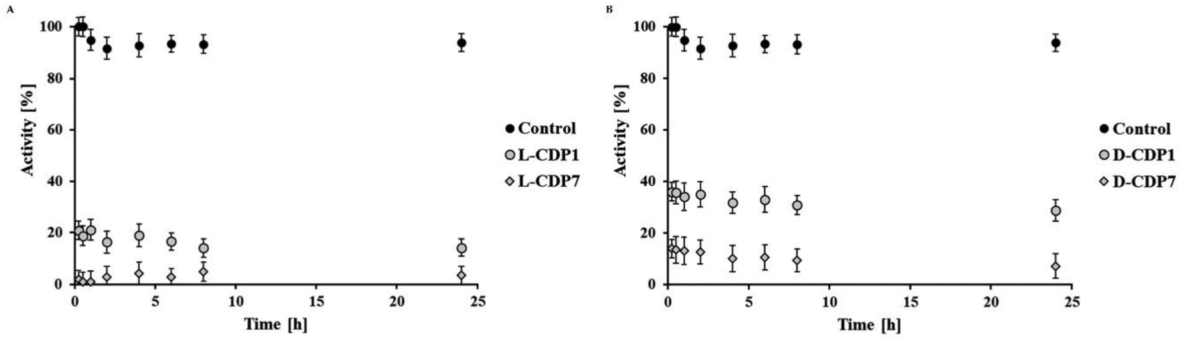


**Figure S11.** 24 h inhibition experiment of L- and D-CDPs against SARS-CoV-2 3CLpro. **A:** 24h inhibition experiment of L-CDP1 and L-CDP7 against SARS-CoV-2 3CLpro. **B:** 24h inhibition experiment of D-CDP1 and D-CDP7 against SARS-CoV-2 3CLpro.

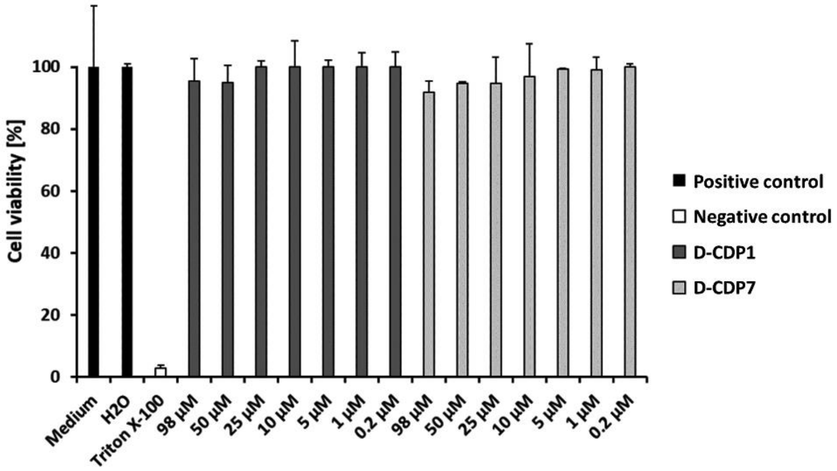


**Figure S12. MTT assay of D-CDP1 and D-CDP7 on Vero CCL-81 cells.** MTT assay was used to evaluate the cytotoxicity of the two D-peptides. Different concentrations of up to 98 µM were used to treat the Vero cells for 2 days. Data shown are the means ± SD from three independent measurements (n=3).

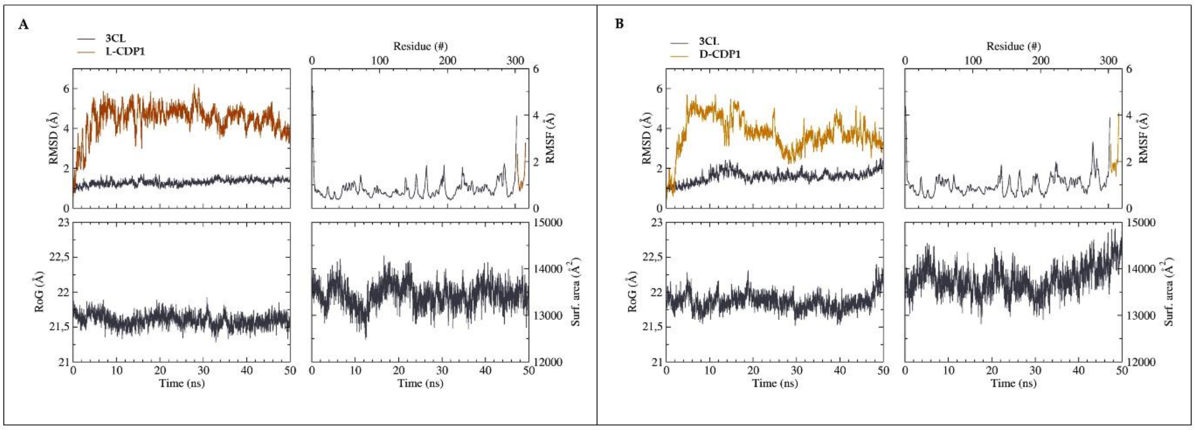


**Figure S13.** Time dependent modifications of the 3CL$^{pro}$/L- D-CDP1 complex. A) 3CL$^{pro}$/L-CDP1 complex. B) 3CLpro/D-CDP1 complex. 3CL$^{pro}$ (gray), L-CDP1 (brown) and D-CDP1 (yellow). RMSD, RMSF, RoG and surface area as function of time. RMSF for each amino acid.

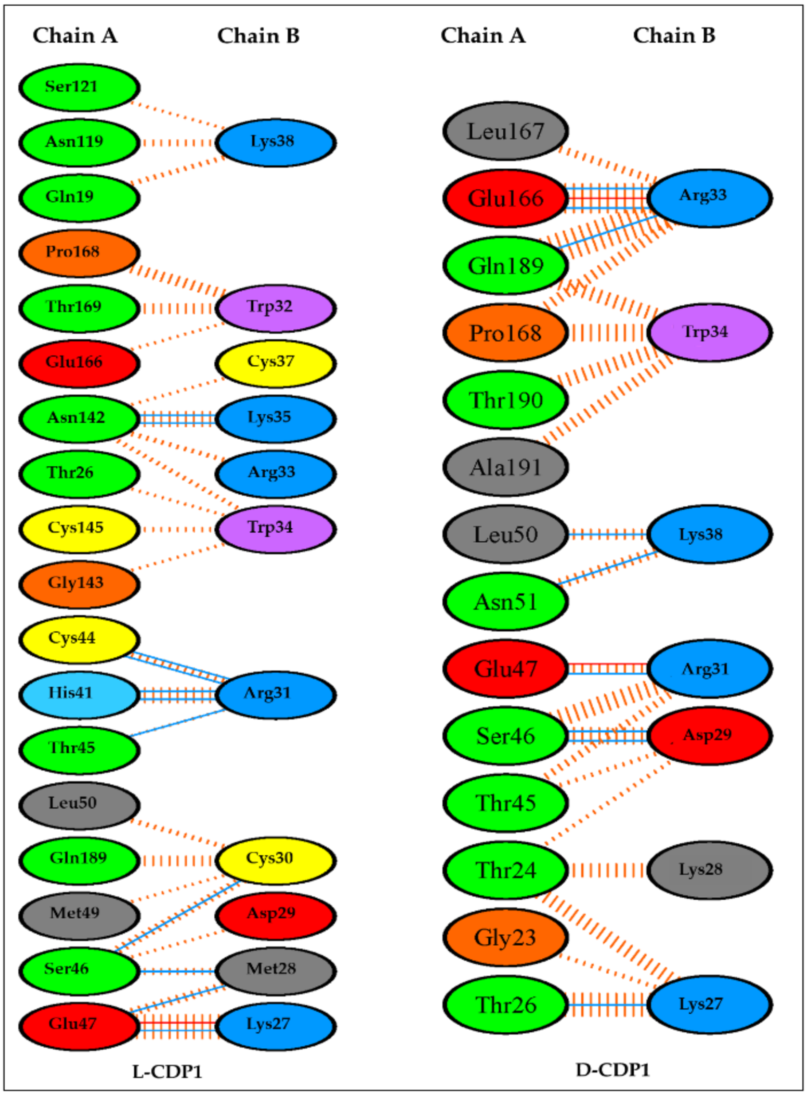


**Figure S14.** List of amino acids- amino acids interactions across 3CLS-peptide interface. The number of H-bond lines between any two residues indicates the number of potential hydrogen bonds between them. For non-bonded contacts, which can be plentiful, the width of the striped line is proportional to the number of atomic contacts. Red line: salt bridges; blue line: hydrogen bonds; orange striped line: non-bonded contacts [2].

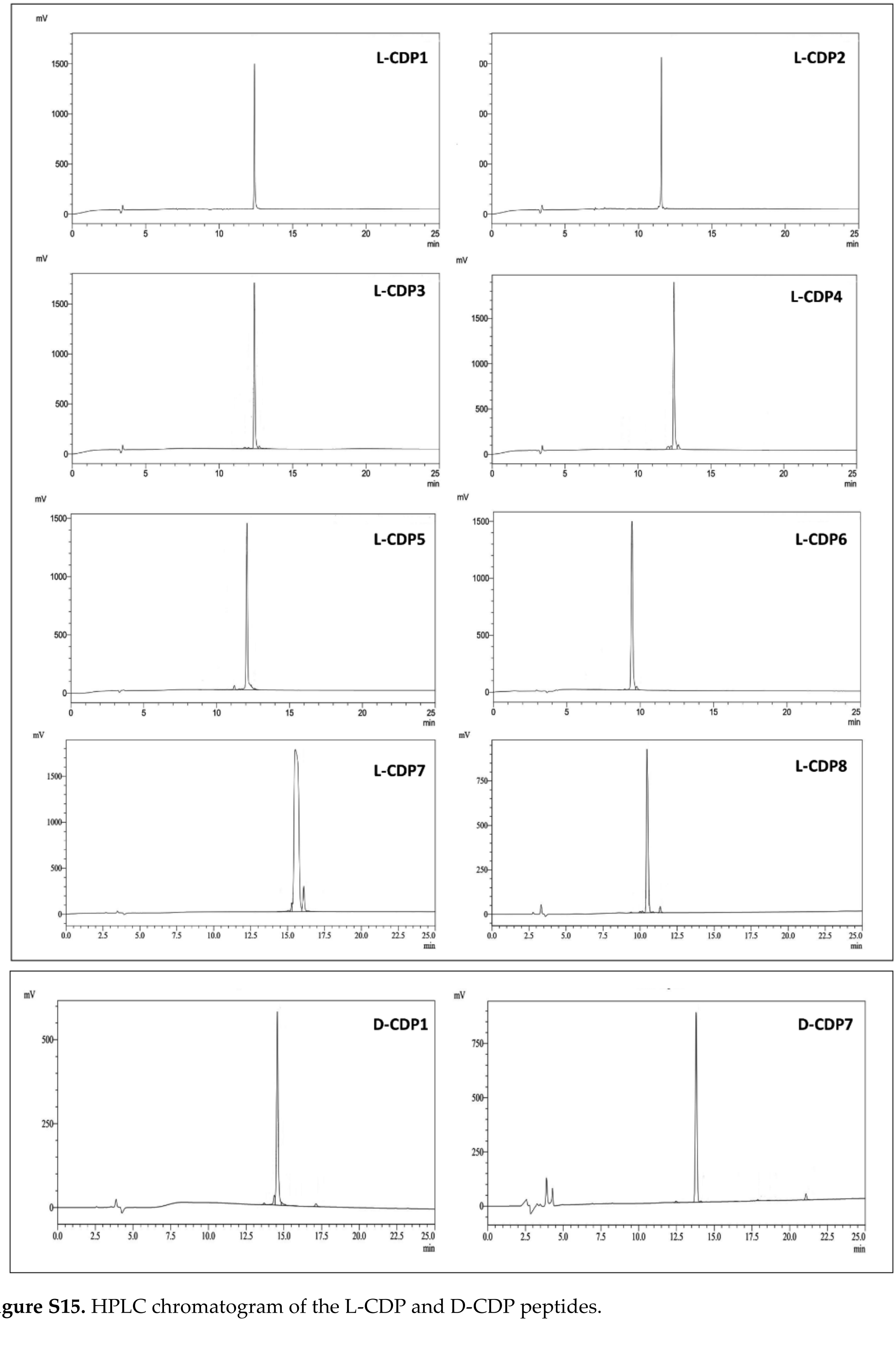


**Figure S15.** HPLC chromatogram of the L-CDP and D-CDP peptides.

**Table S1.** $K_D$ determination of L-CDP1 and L-CDP7 using SPR.

| Peptide | $K_D$1 [nM] | $K_D$2 [nM] | Average $K_D$ [nM] | STD [nM] |
|---|---|---|---|---|
| **L-CDP1** | 89.06 | 40.92 | 64.99 | ± 20.07 |
| **L-CDP7** | 374.3 | 233.7 | 304.0 | ± 70.3 |

**Table S2.** Basic information about tested L- and D-CDPs.

| | Sequence | Conformation | MW | pI | Net charge | Solvent |
|---|---|---|---|---|---|---|
| **CDP1** | KMDCRWRWKCCKK | L and D | 1769.84 | 10.13 | +5 | $H_20$ |
| **CDP2** | KMDSRWRWKSSKK | L | 1721.91 | 11.68 | +5 | $H_20$ |
| **CDP3** | KMDCRWRWKSSKK | L | 1737.89 | 10.99 | +5 | $H_20$ |
| **CDP4** | KMDSRWRWKCCKK | L | 1753.87 | 10.55 | +5 | $H_20$ |
| **CDP5** | KMDSRWRWKSCKK | L | 1737.89 | 10.99 | +5 | $H_20$ |
| **CDP6** | KMDSRWRWKCSKK | L | 1737.89 | 10.99 | +5 | $H_20$ |
| **CDP7** | KMDCRWRWKSCKK | L and D | 1753.87 | 10.55 | +5 | $H_20$ |
| **CDP8** | KMDCRWRWKCSKK | L | 1753.87 | 10.55 | +5 | $H_20$ |
| **CDP9** | RWRWKCCKK | L | 1292.67 | 10.83 | +5 | $H_20$ |

**Table S3.** $K_D$ determination of D-CDP1 and D-CDP7 using MST.

| Peptide | $K_D$1 [nM] | $K_D$2 [nM] | $K_D$3 [nM] | Average $K_D$ [nM] | STD [nM] |
|---|---|---|---|---|---|
| **D-CDP1** | 201.09 | 166.24 | 189.88 | 185.74 | ± 17.8 |
| **D-CDP7** | 1851 | 2012 | 1991 | 1951.3 | ± 87.5 |

**References**


[1] Micsonai, A.; Wien, F.; Bulyáki, E.; Kun, J.; Moussong, E.; Lee, Y.H.; Goto, Y.; Réfrégiers, M.; Kardos, J. BeStSel: a web server for accurate protein secondary structure prediction and fold recognition from the circular dichroism spectra. *Nucl. Acids Res*. 2018, *46*, W315-W322.

[2] Laskowski, RA.; Jabłońska, J.; Pravda L.; Vařeková, RS.; Thornton, JM. PDBsum: Structural summaries of PDB entries. *Prot. Sci.*, 2018, 27, 129-134.
